# Exudate-Guided Janus Trilayer Bioelectronic Dressing for Multiplexed Sensing and Therapy of Chronic Wounds

**DOI:** 10.64898/2025.12.09.692244

**Authors:** Canran Yang, Qiang Wang, Junfeng Yang, Shengwei Shao, Renjie Wang, Chao Luo, Hangwen Li, Jingjie Li, Li Wen

**Affiliations:** Department of Precision Machinery and Precision Instrumentation, University of Science and Technology of China, Hefei, Anhui, 230026, China; The First Affiliated Hospital of USTC, University of Science and Technology of China, Hefei, Anhui, 230001, China; School of Biomedical Engineering, University of Science and Technology of China, Hefei, 230026, China

**Keywords:** trilayer nanofibrous membrane, gradient wettability, antibacterial drug release, electrical stimulation therapy, multifunctional biosensing, in vivo wound healing, wearable bioelectronics

## Abstract

Chronic wound treatment remains challenging because conventional dressings and systemic therapies typically lack real-time adaptability to the dynamic changes in the wound microenvironment. Here, we report a breathable self-adhesive multifunctional wound dressing based on a gradient-wettable Janus trilayer fibrous membrane, which integrates real-time biochemical sensing, on-demand drug release, and low-voltage electrical stimulation therapy simultaneously. The dressing features a vertically integrated trilayer architecture with hierarchical partitioning, where the three vertically stacked layers integrate a multiplex electrochemical sensor array (glucose, lactate, pH), a silver-sulfadiazine drug reservoir, and stimulation electrodes. Specifically, a hydrophilic sensing layer, intermediate drug-storage layer, and hydrophobic adhesive layer are assembled to achieve spatially resolved functionalities. The membrane’s gradient wettability and asymmetric porous structure enable directional exudate transport, thereby triggering stepwise activation of sensing, drug release, and electrical stimulation in a spatiotemporally controlled manner. The dressing is thin, breathable, stretchable, self-adhesive and biocompatible, enabling conformal contact and stable signal acquisition on deformable skin without tapes. This design ensures continuous bio-signal monitoring while maintaining a favorable wound healing microenvironment. Preclinical tests demonstrate its effectiveness in promoting wound closure, reducing infection, and providing continuous wound status monitoring. This integrated system offers a promising approach for next-generation wound care and regenerative medicine.

## 1. Introduction

Chronic wounds impact millions of patients worldwide, causing severe health complications and imposing a substantial economic burden, with an estimated annual healthcare cost of $26.9 billion in China alone^***1–4***^. These wounds, including diabetic foot ulcers, stubborn postsurgical wounds, venous ulcers, and burns, are particularly challenging to treat, as the healing process is often impaired or stalled, accompanied by uncontrolled inflammation and dysfunctional extracellular matrix (ECM) remodeling^***2–6***^. Although existing clinical interventions like skin grafts, skin substitutes, and negative pressure wound therapy can facilitate wound healing, they tend to be invasive, costly, and often inadequate for achieving complete closure^***7***^. Infection is another major obstacle to chronic wound healing, as it can impede tissue regeneration and lead to tissue death or sepsis^(***3***)^. Although local or systemic antibiotics are widely used, their overuse has driven the emergence of antibiotic resistance, making infections harder to control and increasing risks for patients^***8***^. In clinical practice, a variety of conventional wound dressings, including gauze, films, foams, hydrogels, alginates, and silver-impregnated materials, are applied to cover wounds, absorb exudate, and help maintain a moist wound-healing environment. However, these dressings predominantly function as passive barriers, lacking the ability to monitor or dynamically respond to changes in the wound microenvironment. They offer only limited control over local biochemical cues and often fail to inhibit deterioration or reinfection in chronic, highly exuding wounds, which may require frequent and sometimes painful dressing changes^***9–11***^.

To address these limitations, there is growing interest in advanced wound dressings that can dynamically adjust therapeutic intervention to the evolving wound status. Real-time monitoring of wound biochemical parameters is critical to guide therapeutic interventions and facilitate personalized treatment. Wound exudates contain a wide range of metabolites and biomarkers, such as pH, glucose, and lactate, which provide direct insights into the dynamics of tissue repair, inflammation, and infection^***12–15***^. Wearable electrochemical sensors offer a promising approach for continuously converting these biochemical changes at the wound interface into quantitative electrical signals. However, the complex and dynamic composition of wound fluids poses significant challenges to such sensors, particularly in terms of long-term stability, selectivity, and reliable operation in vivo^***13, 16, 17***^. Meanwhile, electrical stimulation has emerged as a promising adjuvant therapy, as it can facilitate fibroblast growth, enhance collagen production, promote keratinocyte migration, and support new blood vessel formation^***18, 19***^. Nevertheless, electrical stimulation alone cannot fully eliminate infection risks, whereas conventional antibiotic delivery often results in uncontrolled dosing and may accelerate the development of antibiotic resistance^***8, 20***^. Therefore, integrating real-time biochemical sensing with synergistic therapies, including localized drug delivery and electrical stimulation, enables dynamic adjustment of treatment regimens in response to evolving wound physiological status, presenting a promising yet largely unrealized strategy for precise, adaptive wound management. These persistent challenges emphasize the urgent need for next-generation wound dressings that transcend passive protection to actively manage the wound microenvironment, tailor therapy dynamically to real-time biochemical cues, and integrate versatile therapeutic functions into a unified multifunctional platform.

Driven by these clinical needs, recent years have witnessed rapid progress in flexible and wearable bioelectronic dressings. A variety of single-layer, planar intergrated systems have been reported, where electrochemical sensors, heaters, or electrical stimulation electrodes are fabricated on elastomer substrates. These systems enable real-time monitoring of temperature, pressure, pH, or key metabolites, and in some cases, can deliver drugs or electrical stimulation directly to the wound site^***14, 21–23***^. However, confining all functional components to a single planar substrate imposes inherent limitations on performance scalability, space utilization, and functional diversity, rendering high-density multifunctional integration extremely challenging. Furthermore, most electronic dressings are fabricated on non- or poorly breathable polymer films with no inherent skin adhesion. Poor breathability causes exudate and sweat accumulation, periwound skin maceration, and wound hypoxia, thereby increasing infection risk and impeding healing. Meanwhile, insufficient inherent skin adhesion necessitates extra fixation tapes or adhesives, resulting in unstable contact during movement and compromising both long-term comfort and stability on irregular, dynamic skin surfaces. Additionally, quantitative and continuous collection of exudate for dynamic wound monitoring poses a significant challenge. In many reported designs^***16, 24, 25***^, exudate must be collected and routed through predefined microchannels to reach the sensing regions. This not only requires precise control over exudate capture and channel geometry but also increases the risk of channel clogging, device failure, and fabrication complexity^***26, 27***^. Ideally, wound dressings should not only meet core clinical requirements, including maintaining a breathable, moist wound microenvironment, preventing secondary infections, and efficiently removing excess wound exudate, but also integrate advanced multifunctional capabilities such as real-time biomarker sensing, controlled drug delivery, and precise therapeutic stimulation all within a single, integrated platform^***14, 20, 28***^. Achieving such comprehensive integration in a structurally simple, mechanically compliant, and clinically practical dressing remains a formidable challenge.

To address above challenges, we developed a multifunctional, wearable bioelectronic dressing platform built on a trilayer Janus fibrous membrane that vertically integrates multiplex electrochemical sensing, localized antimicrobial delivery, and low-voltage electrical stimulation. The trilayer architecture spatially separates distinct functional modules along the thickness direction. The bottom hydrophobic adhesive layer adjacent to the wound integrates an electrical stimulation module. The intermediate layer contains a drug reservoir for controlled release. Meanwhile, the top superhydrophilic layer incorporates a sensing module for continuous detection of multiple biomarkers (glucose, lactate, and pH). Due to the asymmetric gradient wettability of the trilayer membrane, wound exudate can be unidirectionally transported from the wound surface to the outside. During this process, it sequentially triggers the adaptive drug release in the intermediate drug-loaded layer, and initiates the top sensing layer to monitor multiple metabolic biomarkers. The combination of antibiotic drug delivery with electrical stimulation enables substantially accelerated chronic wound closure. Monitoring of multiple biomarkers facilitates timely assessment of wound infection, metabolic, and inflammatory status, thereby providing more comprehensive and personalized information for effective chronic wound management. This innovative design ingeniously converts the directional vertical transport of exudate into a cascading activation mechanism for stacked functional layers. By coupling vertical directional fluid transport with spatially separated functional modules, this bioelectronic dressing enables efficient fluid management, on-demand therapy, and real-time biochemical monitoring simultaneously. Meanwhile, it overcomes the key limitations of conventional single-layer planar integration configuration, thereby achieving high-density integration, device miniaturization, enhanced space utilization, and remarkable functional expandability. Furthermore, this multifunctional electronic dressing exhibits thinness, flexibility, stretchability, breathability, excellent biocompatibility, and self-adhesiveness. It can conformal attachment to diverse body sites and adapt to the dynamic deformations of wound skin. Its superior breathability, unidirectional liquid transport capacity and excellent biocompatibility not only foster an oxygen-permeable microenvironment for tissue regeneration, but also rapidly remove exudate and sweat from the dressing-wound interface, thus effectively preventing infection, inflammation, and skin maceration while ensuring long-term comfort. In addition, its self-adhesiveness enables direct skin attachment, obviating the need for non-breathable auxiliary tapes, and reinforces interlayer bonding to prevent delamination, thus further enhancing both wearing comfort and sensing signal stability. Systematic evaluation in animal models demonstrates that this integrated platform supports multiplex biomarker tracking, significantly accelerates chronic wound healing, and effectively reduces infection and inflammation. This multifunctional bioelectronic dressing offers a promising and convenient all-in-one strategy for wound monitoring and therapy, paving the way for next-generation wound care and regenerative medicine.

## 2. Results and Discussion

### 2.1. Design of the multifunctional gradient-wettable Janus trilayer fibrous dressing

This multifunctional bioelectronic dressing is built on a trilayer electrospun micro-nanofibrous membrane with distinct yet tightly integrated functionalities. It features flexibility, stretchability, breathability, self-adhesion, and gradient wettability, enabling conformal attachment to chronic wound sites with no discomfort or skin irritation, as well as efficient collection and directional transport of small-volume wound exudate. Furthermore, the dressing vertically integrates real-time biochemical sensing, localized drug delivery and low-voltage electrical stimulation, realizing simultaneous wound therapy and dynamic monitoring in a single platform (**Figures 1a, b**). This multifunctional vertical integration relies on the rational design of the trilayer membrane, where each layer is tailored for a specific wound care function. The bottom layer is a hydrophobic adhesive wound-contacting layer composed of thermoplastic polyurethane/pressure-sensitive adhesive (TPU-PSA), which provides robust self-adhesion and mechanical compliance to ensure reliable fixation on wound skin without the need for auxiliary tapes or adhesives. A pair of electrical stimulation electrodes is integrated onto this layer to generate a low-voltage electric field across the wound bed, while the surrounding fibrous layers help distribute deformation and protect the electrodes during bending and stretching. The middle layer is a wettability-transition adhesive layer of thermoplastic polyurethane/polyvinylpyrrolidone/pressure-sensitive adhesive (TPU-PVP-PSA), which allows wettability tuning by adjusting the material ratio and offers reliable adhesion to prevent interlayer delamination. Silver sulfadiazine is loaded within this layer, enabling exudate-mediated controlled hydration and concentration-gradient-driven drug release for localized wound antibacterial and anti-inflammatory therapy. The top layer is a hydrophilic sensing layer of hydrolyzed polyacrylonitrile (HPAN), which is integrated with a flexible sensor array for real-time detection of key biochemical analytes including glucose, lactate, and pH (**Figures 1c, d**). Such a vertical layout separates the electrical stimulation module, drug reservoir, and sensing interface via a trilayer membrane structure, minimizing mutual interference and enabling each module to operate under its optimal local microenvironment.

**Figure 1.**
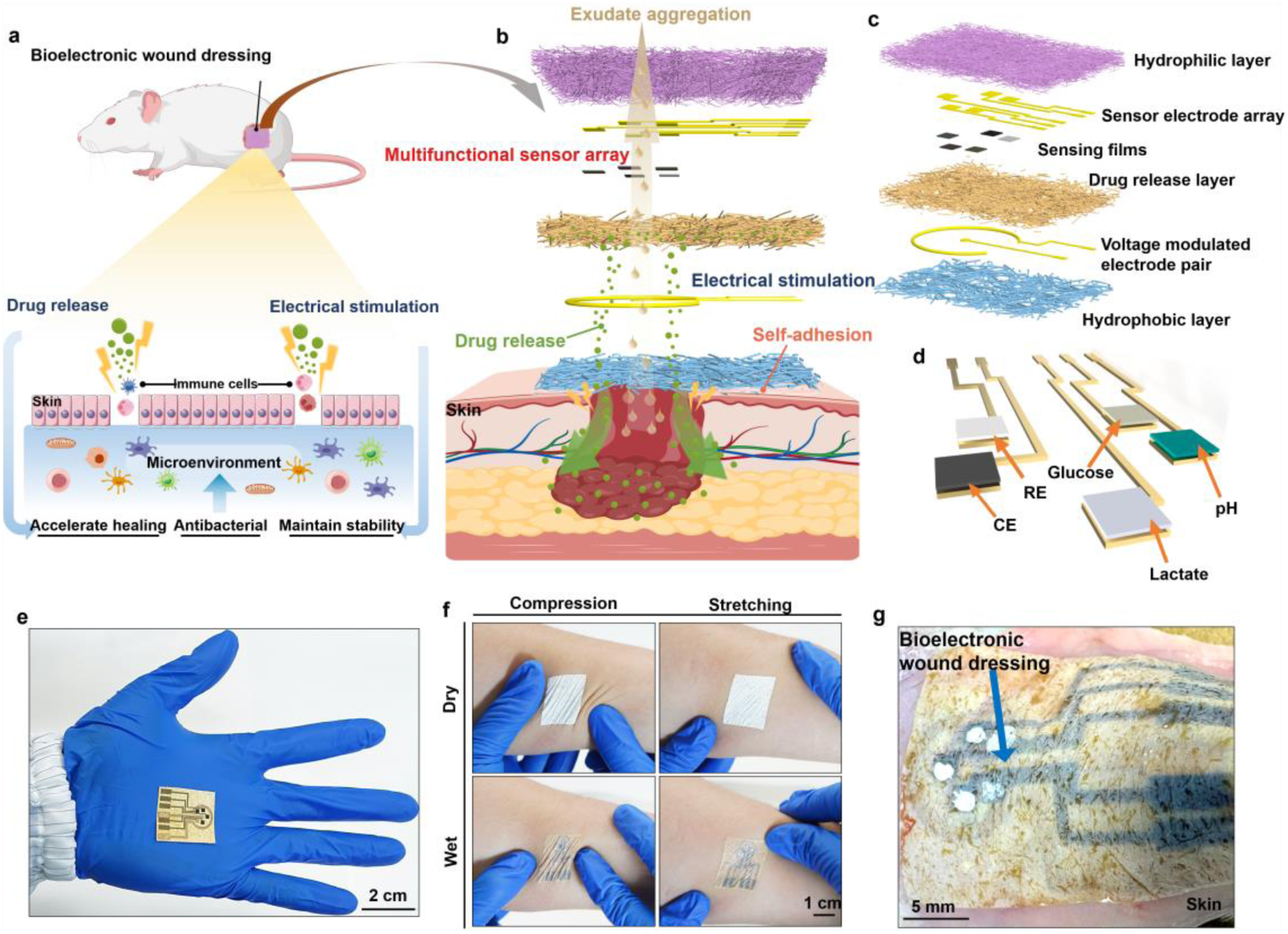
The multifunctional gradient-wettable trilayer bioelectronic dressing for multiplexed sensing and therapy of chronic wounds. **a)** Schematic illustration of the wound-healing mechanism. The dressing accelerates tissue repair by synergistically modulating the wound microenvironment through localized drug release and electrical stimulation. **b)** In situ schematic of the multifunctional dressing. Wound exudate is collected and unidirectionally transported across the stacked trilayer structure to enable simultaneous biochemical monitoring via the multifunctional sensor array, localized antibacterial drug release, and electrical stimulation at the wound interface, with the bottom layer providing self-adhesive fixation to the wound skin. **c)** Exploded schematic of the trilayer architecture, including the hydrophilic sensing layer (sensor electrode array and sensing films), the intermediate drug-release layer, and the hydrophobic self-adhesive contact layer integrated with a voltage-modulated electrode pair. **d)** Schematic layout of the multiplexed electrochemical sensing array (working electrodes for glucose, lactate and pH; RE, reference electrode; CE, counter electrode). **e)** Photographs of the assembled bioelectronic dressing highlighting the integrated electrode layout, including the patterned multiplexed sensing electrodes and the electrical-stimulation electrode pair on the flexible substrate. **f)** Photographs demonstrating conformal self-adhesion and mechanical compliance of the dressing on skin under compression (left) and uniaxial stretching (right) in both dry (top) and wet (bottom) states, indicating stable fixation and shape adaptability under typical skin deformations. **g)** Optical image showing conformal adhesion of the bioelectronic dressing on a murine dorsal wound.

This trilayer membrane exhibits a hydrophobic-to-hydrophilic wettability gradient and a large-to-small pore size gradient from bottom to top. The synergistic effect of these two gradients and the unique capillary action of micro-nanofibers allows pump-free, anti-gravitational transport of wound exudate from the wound bed to the exterior, thus avoiding the need for complex microchannels. (**Figure 1b, c**). Importantly, wound exudate acts not only as the sampling medium but also as a gating stimulus for layer-specific functions. When the exudate front reaches the wettability-transition middle layer, it hydrates the silver sulfadiazine reservoir and initiates concentration-gradient-driven drug release, so antibacterial and anti-inflammatory treatment is enabled only in the presence of exudate. When exudate recedes and the drug layer is no longer hydrated, drug diffusion is suppressed, producing a self-limiting mode of controlled release (**Figure 1b**)^***20, 22***^. As exudate continues to wick into the superhydrophilic HPAN layer, it rapidly spreads to form a well-defined sampling region, activating the sensing films and enabling multiplexed electrochemical monitoring of glucose, lactate, and pH in the chemically complex wound microenvironment (**Figure 1b, d**)^***14, 20, 28***^. Electrical stimulation is realized via a voltage-modulated electrode pair in the TPU-PSA layer closest to the wound interface, providing localized electric field delivery while maintaining continuous exudate transport and sampling through the membrane (**Figure 1c**) ^***17***^.

The overall fabrication process is summarized in **Figure S1** and described in detail in the Experimental Section, with representative intermediate products shown in **Figure S2**. Briefly, a polyacrylonitrile (PAN) membrane is first electrospun and subsequently hydrolyzed to form the HPAN fibrous layer, which serves as the sensing substrate. Gold electrodes for glucose, lactate, and pH sensing are patterned on the HPAN surface, followed by deposition of the corresponding enzyme and pH-responsive films to define the multiplexed sensor array. Next, a TPU-PVP-PSA solution containing silver sulfadiazine is electrospun onto the HPAN sensing layer to form the wettability-transition drug-loaded middle layer, and electrical stimulation electrodes are subsequently sputtered on this layer. After a TPU-PSA solution is electrospun to form the hydrophobic self-adhesive wound-contacting layer, the fully integrated trilayer bioelectronic dressing is finally fabricated (**Figure 1e**). Featuring thinness, softness, breathability, stretchability, and anti-gravitational directional water transport, the dressing conformally adheres to skin reliably in both dry and wet states under compression and stretching. It shows robust self-adhesion and shape adaptability on deformable skin without extra tapes, which is critical for reliable signal acquisition during long-term wear (**Figure 1f**). This vertically integrated multifunctional bioelectronic dressing can conformally self-adhere to the wound surface, providing a compact and extensible platform for on-demand therapy and real-time biochemical monitoring (**Figure 1g**).

### 2.2. Construction and Characterization of the Gradient-wettable Janus Trilayer Membrane

The innovative dressing design employs electrospinning^***28–31***^, a cost-effective and versatile technique that allows precise control over fiber diameter^***32***^ and pore size^***26***^ by adjusting key parameters including polymer solution concentration^***21***^, applied voltage^***33***^, feed rate^***34***^ and deposition time^***35***^. Furthermore, the rational selection of materials with specific physical properties enables the tailoring of distinct membrane characteristics, such as wettability, Young’s modulus, and adhesiveness^***36, 37***^. Based on these considerations, we designed and fabricated a trilayer Janus fibrous membrane with tunable gradient wettability, favorable mechanical strength, and reliable interfacial adhesion via sequential electrospinning, which serves as the structural substrate for the multifunctional bioelectronic dressing.

As illustrated in **Figure 2a**, a PAN solution was first electrospun into a nanofibrous membrane with nanopores, and then hydrolyzed in an alkaline solution to obtain HPAN. This treatment introduced hydrophilic functional groups (–COOH, –CONH) to the fibers, which endowed the HPAN layer with superhydrophilicity (**Figures S3, S4**). Subsequently, a TPU–PVP–PSA solution was electrospun onto HPAN to form the intermediate wettability-transition and adhesive layer. Finally, a TPU–PSA layer was electrospun as the wound-facing contact layer, providing a hydrophobic and strongly adhesive fibrous interface that reinforces overall mechanical integrity and ensures reliable fixation on skin.

**Figure 2.**
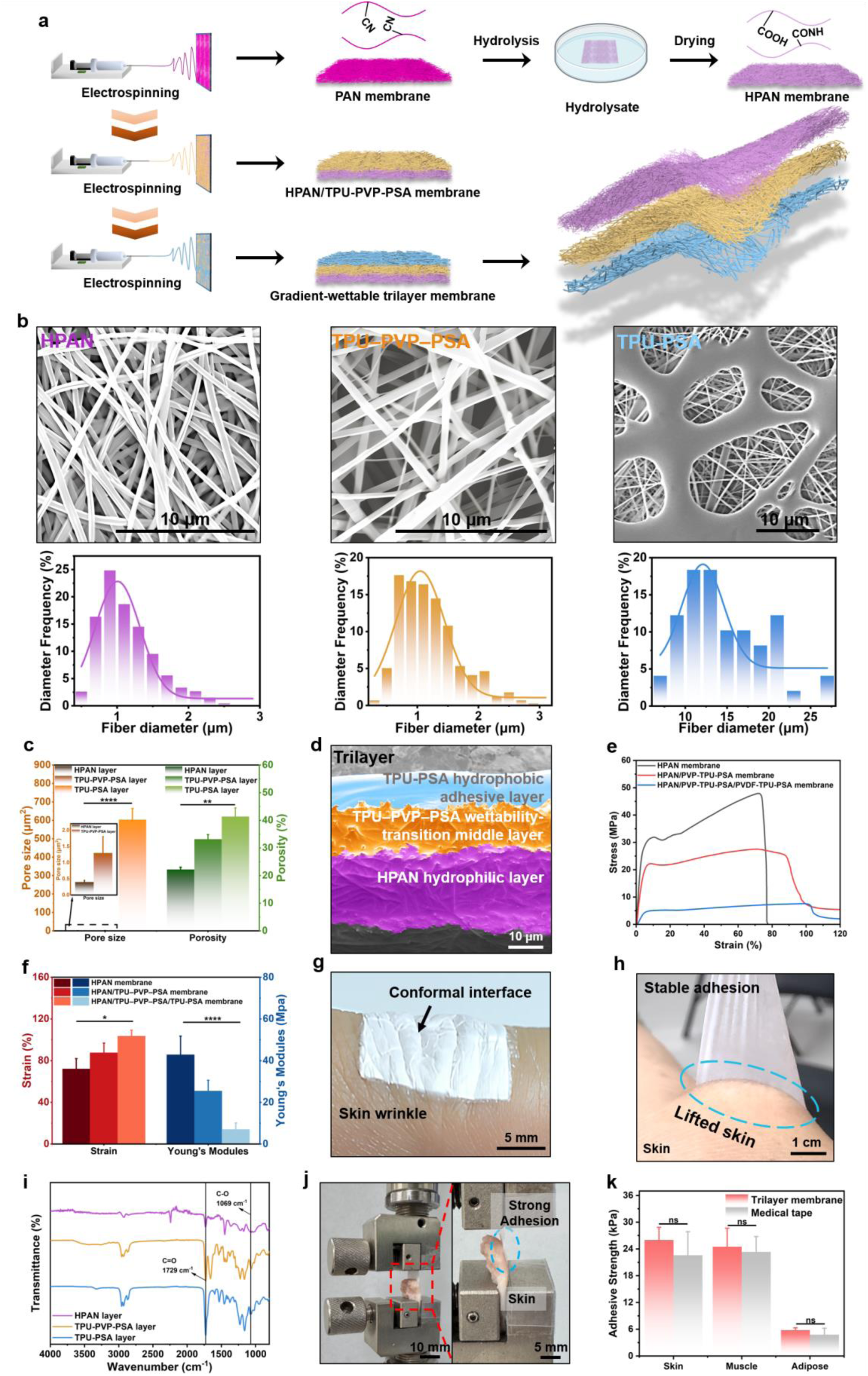
Structural configuration and properties of the gradient-wettable trilayer membrane. **a)** Schematic of the fabrication process involving sequentially electrospinning TPU–PVP–PSA and TPU–PSA layers onto a hydrolyzed PAN nanofibrous membrane (HPAN membrane). **b**) SEM images and corresponding fiber diameter distributions of HPAN layer, TPU–PVP–PSA layer and TPU–PSA layer. **c**) Cross-sectional SEM image of the gradient-wettable trilayer membrane showing distinct layered architecture. HPAN layer (purple), TPU–PVP–PSA layer (orange), and TPU–PSA layer (blue). **d**) Comparison of average pore size and porosity across layers (**P < 0.01; ****P < 0.0001; n = 3). **e**) Representative stress–strain curves of stepwise-assembled monolayer (HPAN layer), bilayer (HPAN and TPU–PVP–PSA layers) and trilayer (HPAN, TPU–PVP–PSA and TPU–PSA layers) membranes. **f**) Comparison of tensile strain and Young’s modulus across stepwise-assembled monolayer, bilayer and trilayer membranes (*P < 0.05; ****P < 0.0001; n = 3). **g)** Conformal adhesion and mechanical compliance of the trilayer membrane on wrinkled skin surfaces. **h**) Optical image showing stable adhesion of the trilayer membrane during peeling from human skin. **i**) FTIR spectra of individual layers showing key functional group peaks that confer distinct adhesion and wettability properties. **j**) Mechanical peel test on porcine skin demonstrating strong adhesion. **k**) Quantitative comparison of adhesion strength on porcine skin, muscle, and adipose tissues (ns p ≥ 0.05; n = 3). Error bars represent mean ± SD.

Morphological analysis of individual layers is presented in **Figure 2b**. SEM images reveal a continuous, bead-free, and porous fibrous network across all three layers. The fiber diameters of the three layers display a gradual increase from 1.0 μm of HPAN layer) to 1.3 μm of TPU-PVP-PSA layerand finally to 14.7 μm of TPU-PSA layer.

This gradient in fiber diameter directly determines the variation in pore size and porosity of the membrane. To quantitatively characterize the porous structure of the multilayer membrane, the average pore size and porosity of each layer were analyzed and are presented in **Figure 2c**. The TPU-PSA layer exhibits the largest pore size (603.6 μm²) and the highest porosity (41.4%), followed by the TPU-PVP-PSA layer (1.3 μm², 33.8%) and HPAN layer (0.4 μm², 22.2%). Such a large-to-small pore size gradient from the TPU-PSA layer to the HPAN layer is crucial for constructing a capillary-driven, bottom-to-top directional fluid transport pathway, which simultaneously ensures efficient exudate management and superior gas permeability. As shown in cross-sectional SEM images ( **Figure 2c**), the trilayer membrane presents distinct interfaces between the top HPAN, intermediate TPU-PVP-PSA, and bottom TPU-PSA layers. Owing to the intrinsic self-adhesiveness of the TPU-PVP-PSA and TPU-PSA layers, the three layers are tightly bonded without delamination. The membrane is ultra-thin and ultra-lightweight, with a thickness of only 48 μm and a low mass of merely 7.1 mg**(Figure S5**), which is promising for imperceptible wear on skin surfaces^***38, 39***^.

The mechanical behavior of the trilayer assembly was evaluated through tensile testing. As shown in **Figure 2e**, the pristine HPAN membrane exhibited limited extensibility (71% elongation at break) and a relatively high tensile strength (47.9 MPa). In contrast, the trilayer architecture delivered a compliant mechanical response, combining enhanced stretchability (101.8% elongation at break) with a reduced tensile strength (7.6 MPa). A quantitative comparison of the maximum tensile strain and Young’s modulus for the layer-by-layer electrospun membranes is presented in **Figure 2f**. The strain at break increased from 72.0% to 103.5%, accompanied by a dramatic reduction in Young’s modulus from 42.8 MPa to 7.0 MPa. This engineered transition from rigid to compliant mechanical behavior is critical for biomedical interfaces, as it significantly mitigates interfacial stress concentration at the skin-device junction. The enhanced compliance minimizes delamination at the membrane-skin interface during movement, thereby improving wear comfort and long-term stability of conformal adhesion for chronic wound management.

Since PSA was incorporated into TPU and TPU-PVP, and both were successfully electrospun into microfibrous membranes, the wound-contacting PSA-TPU layer and the intermediate TPU-PVP-PSA layer exhibit intrinsic adhesiveness. This not only endows the dressing with skin self-adhesion, obviating the need for additional fixation tapes or adhesives, but also strengthens interlayer adhesion between the three layers, preventing interlayer delamination during practical application. As shown in **Figure 2g**, the trilayer membrane can adhere firmly to the wrinkles and fine lines of human skin from the TPU-PSA side. When the membrane was gently peeled from the skin, the underlying skin was visibly lifted upward, indicating strong skin adhesion of the membrane (**Figure 2h**).

The adhesion of the trilayer membrane mainly arises from the intrinsic properties of PSA, a widely used adhesive in commercial tapes. ^***40–42***^. The van der Waals (vdW) forces are the main adhesion sources in PSA^***43***^. Due to the strong distance dependence of vdW forces, stable adhesion relies on the tight conformal contact between the membrane and skin^***44, 45***^. The extremely low Young’s modulus (∼13 kPa) of the membrane facilitates conformal contact with rough and complex skin surface, significantly enhancing the vdW force-based adhesion. In addition to vdW forces, the intermolecular bonds, including hydrogen bonds, covalent bonds, and electrostatic interactions also play an important role in improving the interface adhesion^***46***^. As shown in **Figure 2i**, Fourier-transform infrared (FTIR) spectroscopy of individual layers identifies the key functional groups responsible for the interfacial interactions. The characteristic peaks at ≈ 1729 cm⁻¹ (C=O stretching vibration of urethane/ester groups)^***47, 48***^ and ≈1069 cm⁻¹ (C–O stretching vibration of ether/alcohol moieties)^***49***^ indicate hydrogen-bonding capable sites^***50***^, which can form hydrogen bonds and electrostatic interactions with polar groups (e.g., amino groups and hydroxyl groups) on human skin, thus greatly enhancing adhesion. Notably, the TPU-PSA layer exhibits the highest peak intensities at these positions, followed by TPU-PVP-PSA and HPAN.

This gradient in polar group content directly correlates with adhesion strength. These abundant hydrogen-bonding sites not only further enhance the adhesion between the membrane and skin, but also strengthen interlayer adhesion between the three fibrous layers.

To evaluate the adhesion performance of the trilayer membrane, we quantitatively measured its adhesion strength on different porcine tissues (skin, muscle, and adipose tissue) and compared it with that of commercial medical tape **(Figure 2j)**. As shown in **Figure 2k**, Standard medical tape exhibited adhesive strengths of 22.6, 23.4, and 4.8 kPa when attached to porcine skin, muscle, and adipose tissue, respectively. In comparison, the trilayer membrane achieved corresponding values of 26.0, 24.5, and 5.8 kPa on the same tissue substrates, with no statistically significant difference between the two groups. It is worth noting that although the high oil content of adipose tissue reduces adhesion for both materials, the trilayer membrane still meets the clinical adhesion requirements for skin and muscle applications^***51***^. Importantly, despite its robust adhesive performance, the membrane can be cleanly detached from human skin without leaving adhesive residue or causing epidermal damage (**Figure S6**), which demonstrates its exceptional biocompatibility. This unique combination of strong and safe adhesion, coupled with conformal adaptability to heterogeneous tissue morphologies, ensures secure biomedical device fixation and minimizes the risk of iatrogenic injury during long-term chronic clinical use.

### 2.3. Gradient-wettable Trilayer Membrane for Enhanced Wound Fluid Management and Antimicrobial Delivery

To comprehensively assess the multifunctionality of the gradient-wettable trilayer membrane for wound management, we systematically investigated its gradient wettability, directional fluid transport, and controlled drug release. These integrated features collectively determine its therapeutic efficacy in dynamic wound environments. First, we investigated the differential wettability across the three layers of the membrane. **Figure 3a** illustrates the time-dependent wetting behavior of individual layers and the integrated trilayer membrane via dynamic water contact angle (WCA) measurements using 3 μL droplets. The hydrophilic HPAN layer rapidly transitioned from an initial WCA of 22° to 0° within 9 s, demonstrating its strong hydrophilicity.

**Figure 3.**
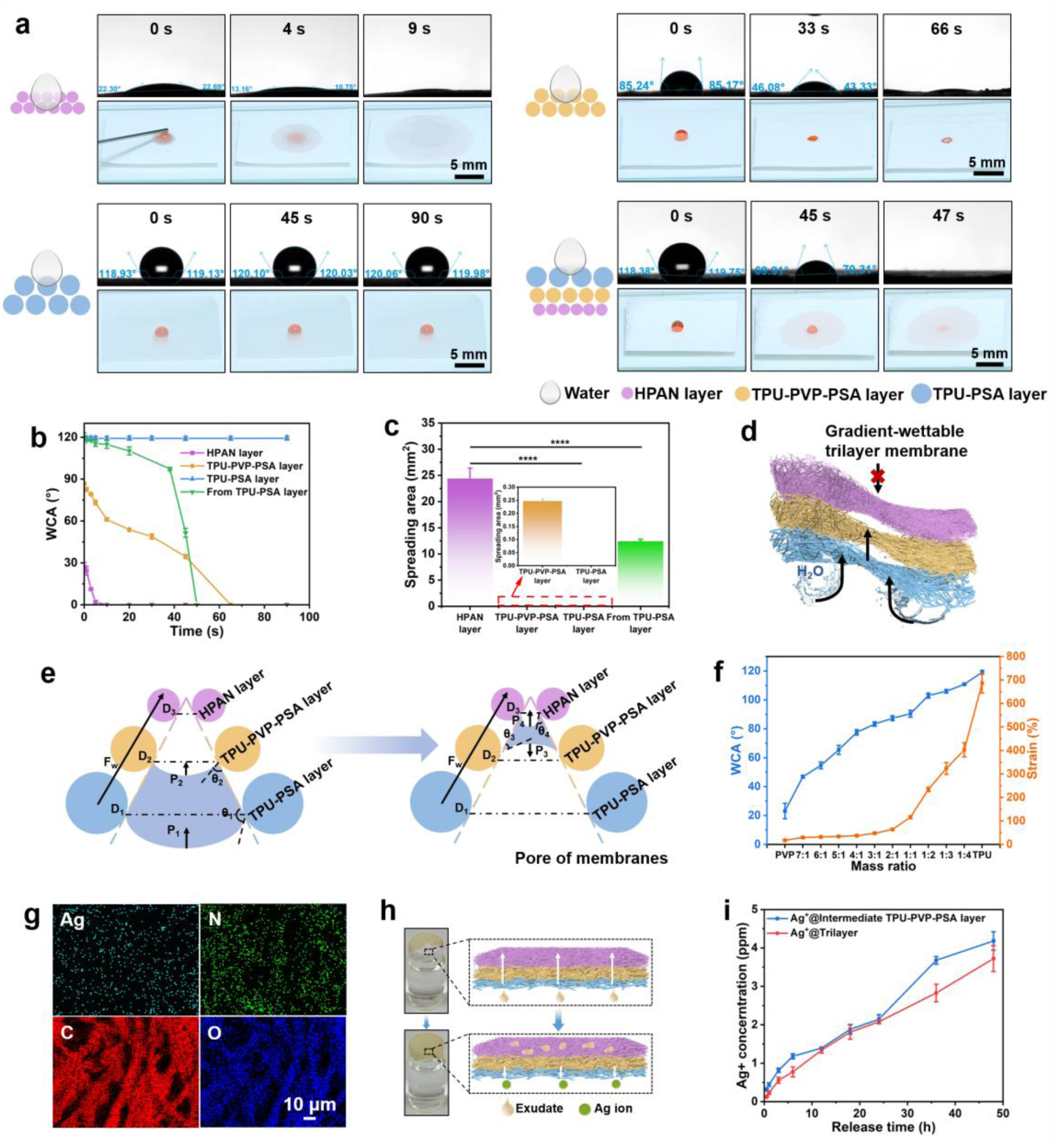
Directional liquid transport and controlled antimicrobial drug release enabled by a gradient-wettable trilayer membrane. **a)** Optical images showing initial 3 μL water droplet contact angles on the hydrophobic adhesive TPU-PSA layer, intermediate wettability-transition TPU-PVP-PSA layer, hydrophilic HPAN layer, and the complete trilayer membrane from TPU-PSA layer. **b)** Water contact angle (WCA) of each individual layer and complete trilayer membrane from hydrophobic TPU-PSA layer at different times (n = 3). **c)** Quantitative analysis of 3 μL water droplet spreading areas after 5 minutes across each individual layer and trilayer membrane (****P < 0.0001, n = 3). **d)** Schematic illustration of the hierarchical trilayer fibrous architecture featuring unidirectional fluid transport and anti-reverse permeation. **e)** The WCA and strain of TPU-PVP-PSA layer at different PVP/TPU mass ratios (n = 3). **f)** Schematic of mechanism underlying unidirectional fluid transport across the trilayer membrane from the hydrophobic TPU-PSA layer to the hydrophilic HPAN layer. **g)** EDS mapping of the TPU-PVP-PSA fibrous membrane. **h)** Schematic and corresponding optical images of unidirectional transport of exudates and directional release of Ag⁺ ions through the trilayer membrane. **i)** Quantitative analysis of Ag⁺ ions cumulative release profiles of from the single wettability-transition intermediate layer and the trilayer membrane over 48 hours (n = 3). Error bars represent mean ± SD.

The WCA of the TPU-PVP-PSA layer decreased from 85° to 0°within 66 s, indicating intermediate wettability. In contrast, the TPU-PSA layer maintained a stable WCA of approximately 120 ° during the measurement, exhibiting stable hydrophobicity. For the full trilayer membrane, when the droplet is dropped from the TPU-PSA side, the WCA decreased from 119° to 0°within 47 s, demonstrating the directional water transport from the hydrophobic side to the hydrophilic side. **Figure 3b** depicts the changes in WCA with time, which clearly demonstrates the distinct wetting dynamics on the different interfaces. Moreover, electrospinning technology endows the trilayer membrane with an intrinsic porous network structure, resulting in natural breathability. **As shown in Figure S7**, the membrane delivers superior moisture vapor transmission rates (MVTR) of 5.7 kg·m⁻²·d⁻¹, significantly higher than that of conventional PDMS substrates (0.5 kg·m⁻²·d⁻¹) ^***22***^ at 50 °C and 30% RH. By preventing fluid accumulation while maintaining favorable oxygen permeability, the membrane avoids the formation of hypoxic microenvironments that impede angiogenesis and epithelial regeneration^***52***^.

To complement the dynamic wetting analysis, 3 μL water droplets were dropped separately onto each individual layer (HPAN, TPU-PVP-PSA and TPU-PSA) and onto the TPU-PSA side of the complete trilayer membrane (**Figure S8**). Quantitative analysis of the spreading areas in **Figure 3c** reveals that the HPAN layer achieved the largest spreading area (24.41 mm²), followed by the complete trilayer membrane (12.39 mm²). In contrast, the TPU-PVP-PSA layer showed only limited spreading (0.25 mm²), while the TPU-PSA layer exhibited negligible wetting (0 mm²). The gradient wettability of the trilayer membrane endows it with asymmetric liquid handling capability, enabling unidirectional water transport from the bottom hydrophobic layer to the upper hydrophilic layer. This design not only ensures efficient exudate management via unidirectional drainage from the wound-contacting hydrophobic layer to the outermost hydrophilic layer, but also effectively blocks reverse water permeation and retrograde contaminant ingress (**Figure 3d**).

The antigravity directional water transport is governed by both the capillary pressure gradient (Laplace pressure) within the interconnected micro/nano fibrous pores and surface energy gradient force (*F_W_*) induced by wettability gradient (**Figure 3e** and **Figure S9**). When a water droplet contacts the TPU-PSA layer of the trilayer membrane from below, it penetrates into the interfibrillar capillary channels under the upward surface energy gradient force *F_W_* and forms a curved meniscus that generates capillary pressure, which is expressed as^***53***^:

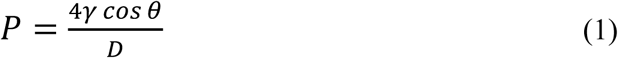

where γ is the liquid–gas interfacial tension, θ is the contact angle on the fiber surface, and *D* is the pore diameter. Because the trilayer membrane integrates a wettability gradient from the hydrophobic TPU-PSA side to the superhydrophilic HPAN side, along with a pore size gradient from larger to smaller pores, capillary pressure *P* increases upward across the membrane thickness, producing a net pressure bias that drives fluid unidirectionally upward against gravity. In parallel, the surface energy gradient force *F_W_* at the layer interfaces favors spreading and wetting toward the more hydrophilic region, further promoting upward wicking through the laminate. Furthermore, the hydrophobic TPU-PSA contact layer can prevent inward wetting of externally deposited aqueous droplets and suspended contaminants (including bacteria and particulates), thus reducing the the risks of exudate accumulation, tissue maceration, and secondary infection.

It is noted that the wettability and mechanical compliance of the TPU–PVP–PSA intermediate layer can be regulated by adjusting the mixing ratio of TPU and PVP (**Figure 3f**). As the TPU content increases, the WCA increases from about 23° to 120°, indicating a progressive shift toward a more hydrophobic surface. Meanwhile, the tensile strain at break rises sharply from about 17.5% to 686%, reflecting a remarkable improvement in stretchability. This trend is consistent with the inherent hydrophobic and elastomeric characteristics of TPU, which reduces surface polarity while endowing the network with enhanced chain mobility and elastic extensibility. In contrast, the hydrophilic PVP facilitates surface wetting yet tends to form a stiffer and less stretchable network. Such tunability enables precise compositional tailoring of the intermediate-layer to meet practical requirements, achieving an optimal balance between fluid-transport performance and mechanical robustness. To achieve both graded wettability and reduced Young’s modulus in the trilayer system, a 1:1 TPU-to-PVP ratio was selected. This formulation endows the intermediate-layer with a WCA of 85.20° and an elongation at break of 115.73%. Thus, it acts as a bridge between the hydrophilic outer layer and hydrophobic inner layer, enabling the trilayer membrane to achieve continuous directional water transport while maintaining mechanical compliance.

Beyond effective fluid management, the incorporation of antimicrobial components is essential for preventing wound infections^***20, 26***^. In this study, silver sulfadiazine was incorporated into the intermediate TPU-PVP-PSA layer. Energy-dispersive X-ray spectroscopy (EDS) mapping of this layer (**Figure 3g**) demonstrates that Ag, C, N and O elements are homogeneously distributed throughout the fibrous network, indicating uniform loading of silver sulfadiazine without obvious aggregation. When wound exudate permeates this wettability-transition region, Ag⁺ ions from silver sulfadiazine diffuse toward the wound bed along a concentration gradient^***54, 55***^. **Figure 3h** schematically illustrates this directional ion release toward the wound interface. Notably, the gradient-wettability architecture spatially separates fluid and ion transport pathways. Wound exudate is rapidly wicked upward via the combined effects of capillary action and gradient wettability, which not only prevents fluid accumulation on the wound surface but also hydrates the TPU–PVP–PSA intermediate layer and dissolves silver sulfadiazine, thereby triggering in situ Ag⁺ generation. The released Ag⁺ ions then migrate downward toward the wound interface through the interconnected fibrous network, driven primarily by concentration-gradient diffusion. **Figure 3i** presents a quantitative analysis of Ag⁺ release from the intermediate single-layer and the trilayer membrane in 10 mL of 0.1× PBS. The intermediate single layer and the trilayer membrane released 4.2 ppm and 3.7 ppm of Ag⁺ within 48 h, respectively. Both concentrations are well above the bactericidal threshold of 0.1 ppm, thus meeting the therapeutic requirement^***56, 57***^. This indicates that drug release driven by the concentration gradient is unaffected by the wettability of the trilayer membrane, enabling sustained and effective antibacterial activity while ensuring efficient exudate management. By coupling directional fluid drainage with sustained silver ion release, the membrane establishes a stable, infection-resistant microenvironment conducive to tissue repair.

### 2.4. Design and characterization of a multiplexed sensor array for biomarker analysis and electrical stimulation

Driven by the pore-size gradient and wetting gradient of the trilayer fibrous membrane, the wound exudate is further transported upward to the top hydrophilic layer, where the integrated biosensing array is triggered to enable real-time multiplexed monitoring of the biomarkers including glucose, lactate, and pH in wound exudate. Continuous and selective measurement of these key biomarkers is particularly critical for assessing the wound healing status^***14, 58***^. In the sensing array, glucose and lactate are detected by amperometric enzymatic electrodes, in which glucose oxidase (GOx) and lactate oxidase (LOx) catalyze the oxidation of their corresponding substrate to generate hydrogen peroxide or reduced mediator species. At a fixed applied potential, these redox processes generate a Faradaic current proportional to the local analyte concentration. The pH is monitored by a potentiometric electrode coated with a polyaniline-based sensing film. The protonation state varies with the hydrogen ion activity at the interface, generating a potential with a nearly linear dependence on pH (**Figures 4a-c**). To achieve stable and selective operation in the chemically complex wound exudate environment, the sensing interfaces are further engineered at the material level. Tetrathiafulvalene (TTF) combined with carbon nanotubes (CNTs) forms a conductive composite membrane that efficiently immobilizes GOx and LOx and facilitates rapid electron transfer, thus enhancing sensitivity, response rate, and operational stability^***59***^. For the glucose sensor, GOx is further immobilized in a bovine serum albumin (BSA) and Nafion matrix which serves as an ion diffusion limiting layer, thereby improving sensitivity and stability while suppressing interference from electroactive species^***60***^. Given that lactate faster diffusion owing to its smaller molecular size, chitosan is introduced to enhance enzyme immobilization and interfacial stability, accelerate electron transfer, and reduce interference via electrostatic repulsion. Meanwhile, a Polyvinyl Chloride (PVC)-based membrane provides a denser diffusion-limiting barrier to compensate for the faster mass transport of lactate and extend the linear range^***61***^. These tailored interfacial structures are integrated on the top HPAN layer of the trilayer membrane, where gradient wettability ensures continuous wicking of wound exudate toward the sensing surface and replenished by fresh fluid, effectively enhancing analyte accessibility and sampling efficiency.

**Figure 4.**
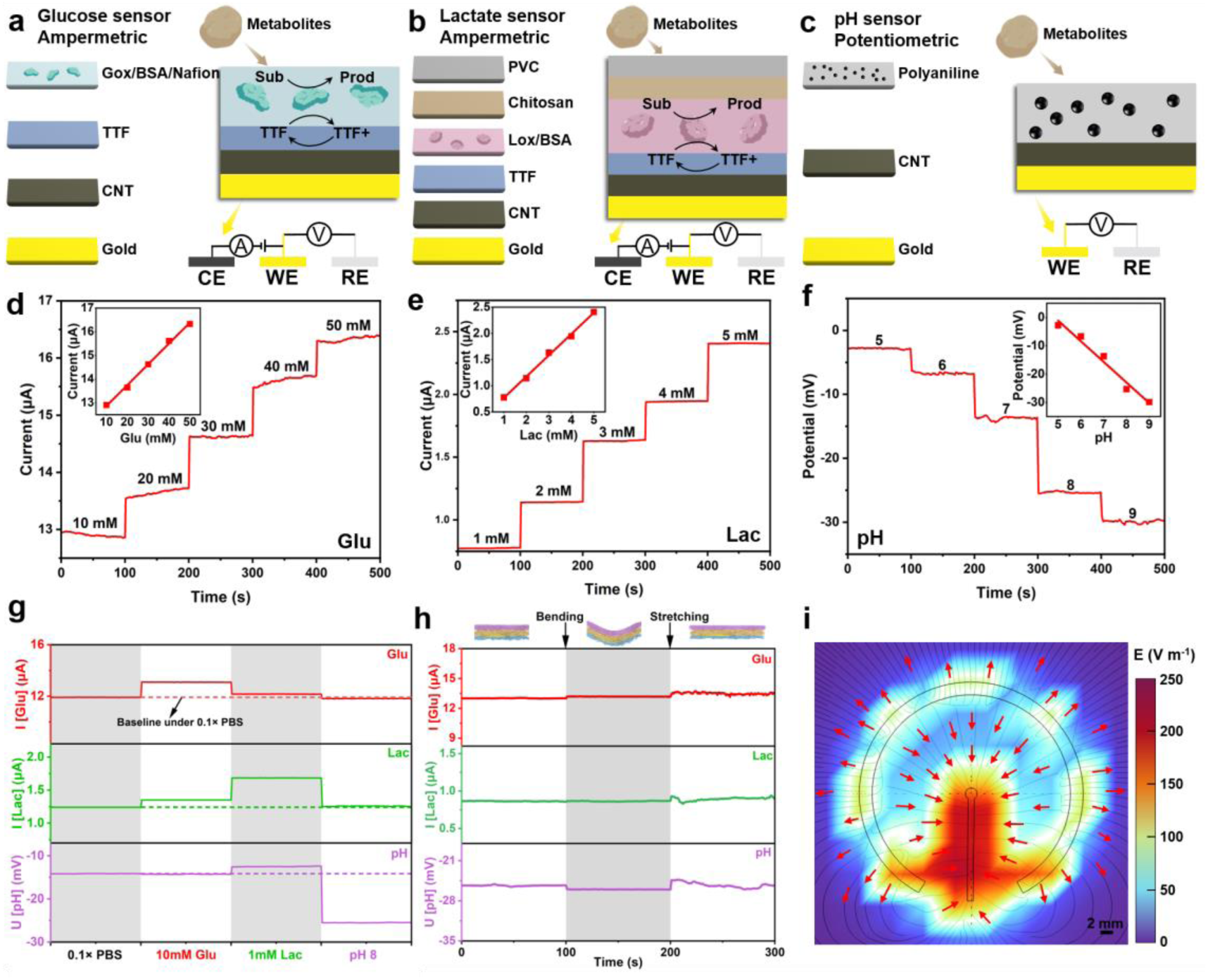
Characterization of the multiplexed electrochemical sensing array and electrical stimulation electrodes in the trilayer bioelectronic dressing. a-c) Schematics of **(a)** glucose sensor, **(b)** lactate sensor, and **(c)** pH sensor. Gox, glucose oxidase; BSA, Bovine Serum Albumin; TTF, tetrathiafulvalene; CNT, carbon nanotube; Sub, substrate; Prod, product; PVC, polyvinyl chloride; Lox, lactate oxidase; CE, counter electrode; WE, working electrode; RE, reference electrode. **d-f)** Chronoamperometric responses of the enzymatic **(d)** glucose sensor, **(e)** lactate sensor, and potentiometric response of the **(f)** pH sensor. Insets show the corresponding calibration plots with linear fits. **g)** Selectivity study of the sensors. Time-dependent responses recorded in separate measurements where 50 μL of 0.1× PBS, 10 mM glucose, 1 mM lactate, and pH 8 standard buffer was individually added onto the sensors; each curve corresponds to one test solution. **h)** Responses of the multiplexed sensor array before and during mechanical bending (radius 1 cm) and mechanical stretching (10% strain) in 0.1× PBS (pH 7) containing 10 mM glucose and 1 mM lactate. **i)** Numerical simulation of the electrical field generated by the custom-designed electrical stimulation electrodes during operation. E, electrical field.

The electrochemical performance of the glucose, lactate, and pH sensors are summarized in **Figures 4d-f**. Both the glucose and lactate sensors exhibit monotonic increases in steady-state current with increasing respective analyte concentrations. They exhibit excellent linear calibration responses over the tested ranges, with correlation coefficients of 0.998 for both sensors, and sensitivities of 0.088 μA mM⁻¹ for glucose and 0.406 μA mM⁻¹ for lactate. The pH sensor displays a nearly linear potential response to solution pH, with a correlation coefficient of −0.986 and a sensitivity of −7.278 mV pH⁻¹, which is sufficient for monitoring relative pH variations. All three sensors employ Ag/AgCl coated electrodes as reference electrodes to maintain a stable potential^***62***^. The high linearity and sensitivities of all three channels indicate that the sensor array can reliably distinguish subtle variations in glucose, lactate and pH. Combined with the conformal, self-adhesive and breathable trilayer membrane, these sensing characteristics establish a robust basis for multiplexed biochemical monitoring directly at the wound interface.

Wound exudate contains a complex mixture of metabolites that may potentially interfere with sensor readings. To evaluate the selectivity of the three sensors, their responses to both target and non-target metabolites were tested. As shown in **Figure 4g**, all three sensors showed negligible responses to non-target metabolites, while generating significant signals toward their target analytes. Notably, short-term fluctuations in environmental pH had almost no effect on the glucose and lactate sensor responses (**Figure S10**). The operational stability of the three sensors was further assessed by repeated measurements. In these tests, 5 μL aliquots of test solution were repeatedly dropped onto the sensing area, measured, and then replaced with fresh solution in successive cycles. As shown in **Figure S11**, the pH sensor exhibited only minor response variations before and after repeated operation, and likewise the signals from the glucose and lactate sensors remained nearly unchanged, demonstrating excellent stability and reproducibility during prolonged use. Furthermore, the electrospun trilayer fibrous membrane is intrinsically soft and stretchable, allowing conformal adhesion to the skin and withstanding common deformations including bending and tension. To verify its mechanical robustness under practical wearing conditions, we monitored the sensor outputs during controlled bending and uniaxial stretching. Mechanical bending and 10% uniaxial stretching (**Figure 4h**), as well as repective bending cycles (**Figure S12**) caused no significant changes in the responses of all three sensors, further verifying the mechanical reliability and stability of the system.

In addition to multiplexed biosensing capabilities, the electronic dressing is integrated with an electrical stimulation module, which produces a directional electric field to promote tissue regeneration and accelerate wound healing. The directional electric field is generated by a pair of custom-designed stimulation electrodes, which can be precisely regulated via voltage modulation (**Figures S13 and S14**). **Figure 4i** presents a numerical simulation of the directional electrical field. The simulation results show a directional distribution of the electric field across the wound surface, which provides effectiveness electrical stimulation to regulate cellular behaviors including cell-cell junctions, cell migration, oriented cell division, and related processes, thereby enhancing tissue regenerative and accelerating wound healing^***19, 63–66***^.

### 2.5. In vivo assessment of the multifunctional wound dressing in a murine model

To verify the performance and effectiveness of our electronic dressing, in vivo preclinical assessment is essential. Firstly, we evaluated the biocompatibility of the trilayer membrane to verify its biosafety for biomedical applications. NIH-3T3 fibroblasts were co-cultured with the membrane extracts to examine cytocompatibility (**Figure 5a**). After 48 h of incubation, cells from the three groups (control, drug-free extract, and drug-loaded extract) were subjected to live/dead staining and imaged via fluorescence microscopy (**Figure 5b**). As expected, only a small number of dead cells (red) were observed among the abundant live cells (green) in all three groups, with no significant differences in cell morphology and viability detected among the groups. To further quantify cell viability, optical density was determined using the Cell Counting Kit-8 (CCK-8) assay. As shown in **Figure 5c**, no significant differences were observed among the groups over the 48 h culture period, indicating that the drug-loaded membrane exhibited no cytotoxicity. These results demonstrate that the membrane imposes no adverse effects on cell viability and is suitable for subsequent in vivo investigation.

**Figure 5.**
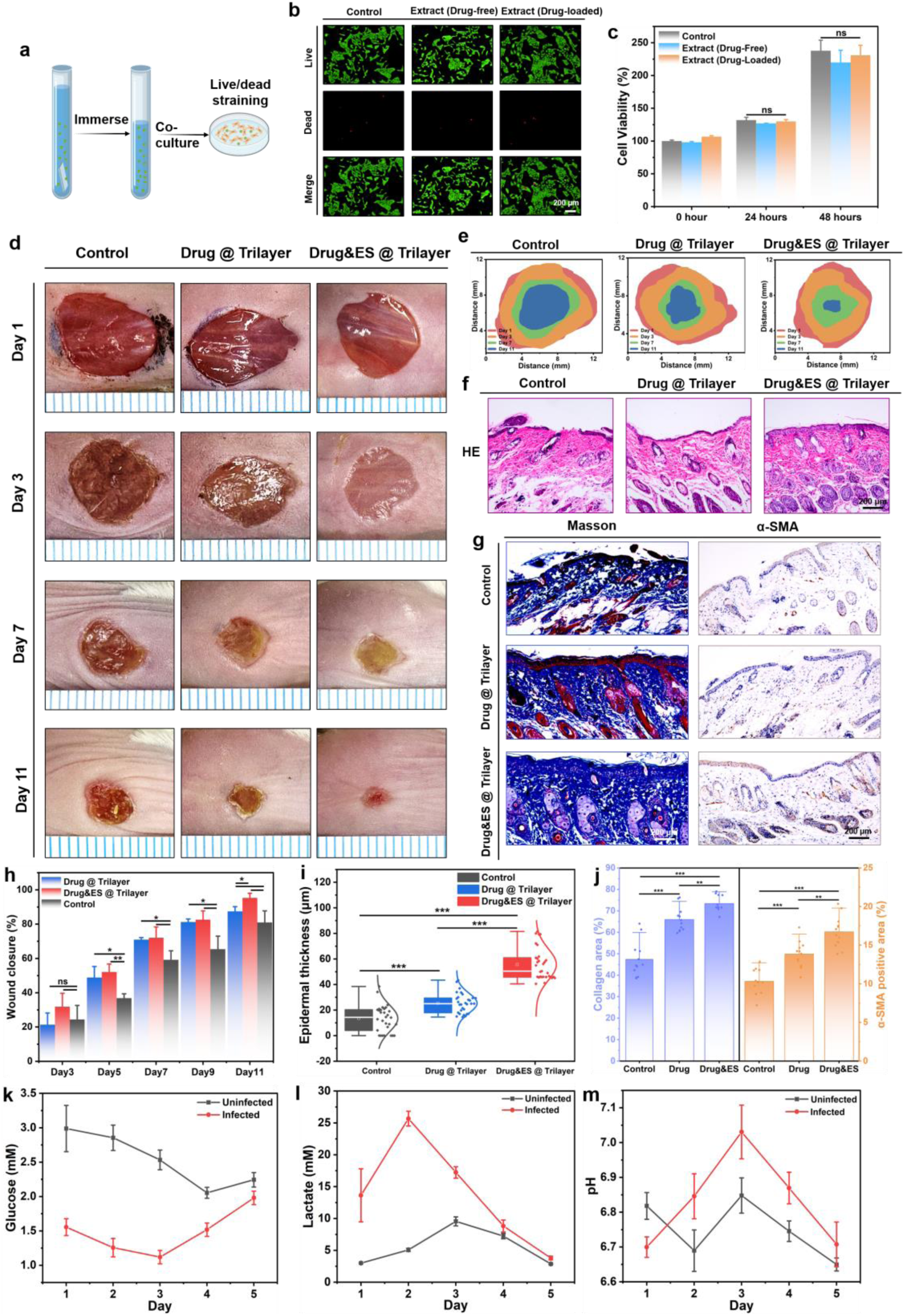
Preliminary in vitro and in vivo evaluation of the trilayer wound dressing in rats. **a)** Schematic illustration of the preparation of the membrane extract used for live/dead staining. **b)** Live/dead fluorescence images of the control group, extract (drug-free), and extract (drug-loaded) after 48 hours of culture. **c)** CCK-8 cytotoxicity test of the various groups at the time points of 0, 24, and 48 hours (n = 3). **d)** The representative photographs of wound healing with different treatment groups on days 1, 3, 7, and 11. **e)** The wound trace of these groups. **f)** HE staining images of wound samples. **g)** Staining of Masson and α-SMA in various treatment groups. **h)** Statistical results of wound closure rates on days 3, 5, 7, 9, and 11 (*P < 0.05; **P < 0.01; n = 3). **i)** The epidermal thickness of repair areas in different treatment groups (**P < 0.01; ***P < 0.001; n = 30). **j)** Expression area of collagen (Masson) and α-SMA in various treatment groups (**P < 0.01; ***P < 0.001; n = 12). **k-m)** Continuous monitoring of the **(k)** glucose, (**l)** lactate, and (**m)** pH of the infected/uninfected wounds during the healing process (n = 3). Error bars represent mean ± SD.

To evaluate its efficacy in combined therapy and wound healing, we applied the bioelectronic trilayer dressing to a full-thickness dorsal skin defect model in rats (**Figure 5d**). Three groups were included: an untreated control group, a drug-only dressing group, and a drug loaded dressing combined with electrical stimulation group (drug & ES). Benefiting from its ultrathin, flexible, and stretchable fibrous structure as well as intrinsic self-adhesion, the dressing can conformally adhere to the wound site without additional tapes, ensuring stable contact during animal movement. Compared with the control group, the drug-loaded trilayer dressing markedly accelerated wound contraction and epithelialization. This therapeutic effect can be attributed to localized antimicrobial and anti-inflammatory treatment, which helps reduce bacterial burden and modulate the immune response at the wound site. The addition of electrical stimulation further amplified the therapeutic effect, consistent with the well-established role of exogenous electric fields in promoting the migration and proliferation of keratinocytes and fibroblasts. After 11 days of treatment, the drug & ES dressing achieved a wound closure rate of 95.27%, outperforming both the drug-only dressing (87.48%) and the control group (80.96%) (**Figures 5e, h**, and **Figures S15-S17**). Hematoxylin–eosin (HE) staining of the regenerated tissue (**Figure 5f**) confirmed these macroscopic observations. In the drug & ES dressing group, the wound site was covered by a continuous, stratified epidermal layer, along with a well-organized dermal structure and dense granulation tissue. By contrast, the drug-only dressing and control groups still exhibited residual scabs, surface edema and discontinuous or thinner epidermis. Quantitative analysis showed that the epidermis is substantially thicker in the drug & ES dressing group (55.88 μm) than in the drug-only dressing group (26.18 μm) and the control group (12.95 μm) (**Figure 5i**), indicating more advanced re-epithelialization and tissue remodeling under the combined drug and electrical stimulation treatment. To further evaluate collagen deposition, tissue maturation, and myofibroblast activation in regenerated tissue, Masson trichrome and α-SMA immunohistochemical staining were performed on wound samples from different treatment groups **(Figure 5g)**. The results showed that wounds treated with the drug & ES dressing group exhibited the highest collagen area (73.38%), significantly higher than that in the drug-only dressing group (65.94%) and the untreated control group (47.35%). Similarly, the α-SMA-positive area, an indicator of myofibroblast presence and tissue remodeling, was also the highest in the drug & ES dressing group (16.72%), followed by the drug-only dressing group (13.85%) and the control group (10.32%) (**Figure 5j**). These findings suggest that combined drug and electrical stimulation not only promotes collagen deposition but also enhances myofibroblast activation, thereby accelerating tissue remodeling and repair during the early healing stage.

Meanwhile, the integrated sensing module on the electronic dressing enables real-time, continuous monitoring of wound biochemical markers, providing dynamic information that complements the end-point histology and closure metrics. To investigate the dynamics of wound-related biochemical parameters under infected and uninfected conditions, we established two full-thickness skin wound models with distinct microenvironments. In the infected model, wounds were inoculated with Staphylococcus aureus to induce sustained inflammatory and infectious burden, whereas the uninfected model followed a normal healing trajectory without deliberate microbial challenge. We continuously monitored glucose and lactate concentrations as well as the pH levels in both wound models (**Figures 5k-m** and **Figures S18-S21**). Throughout the 5-day observation period, infected and uninfected wounds exhibited distinct profiles in glucose, lactate, and pH levels. Glucose levels were consistently lower in infected wounds, attributable to increased local consumption by infiltrating immune cells and bacteria, as well as impaired perfusion in inflamed tissue^***67***^. Lactate levels were higher in infected wounds, consistent with enhanced glycolytic metabolism under inflammatory stress and limited oxygen transport in the infected microenvironment^***68***^. Wound pH also shifted upward in infected wounds, reflecting the transition toward a more alkaline milieu commonly associated with bacterial metabolic byproducts, protease activation, and hypoxia-driven inflammation^***69, 70***^. Overall, these trends are consistent with previous reports and validate the ability of the integrated dressing for monitoring infection-associated biochemical signatures in situ. In the uninfected group, transient elevations in lactate and pH during the early post-injury period were observed, which can be reasonably attributed to the acute inflammatory phase together with temporary, local hypoxia caused by vascular disruption immediately after injury, followed by gradual normalization as perfusion and tissue remodeling proceed. Notably, the peaks of lactate and pH typically appeared on postoperative day 2 or 3, corresponding to the inflammatory phase during which large numbers of leukocytes infiltrate the wound site^***2, 71***^. After this period, both parameters exhibited a gradual decline in infected and uninfected wounds. Although the infected wounds maintained relatively higher levels, the continuous downward trend indicates progressive healing, likely attributed to the antibacterial activity of the trilayer dressing. The above in vivo experimental results demonstrate that the combined drug and electrical stimulation therapy integrated within the electronic dressing can effectively promote tissue regeneration and accelerate wound healing. Additionally, continuous dynamic monitoring of key biochemical indicators in wound exudate enables timely adjustment and optimization of treatment regimens according on the actual wound status, thereby paving the way for monitoring-guided personalized wound management.

## 3. Discussion

In this work, we developed a fully integrated wound dressing that incorporates biosensing, electrical stimulation and controlled drug release into a trilayer asymmetric fibrous membrane. This system enables comprehensive, continuous monitoring of wound biochemical indicators while simultaneously accelerating tissue repair and wound healing. The soft, stretchable, breathable, self-adhesive, and biocompatible trilayer fibrous membrane provides a reliable platform for integrating of multiple functional modules, while ensuring long-term wearing comfort and stability. The asymmetric wettability structure effectively drains wound exudate from the wound bed to the outer layer, preventing excessive moisture, skin maceration and inflammation. During the vertical upward fluid transport, wound exudate infiltrates the drug-loaded intermediate layer. The resulting concentration gradient drives the diffusion of antibacterial agents toward the wound site, exerting effective antimicrobial and anti-inflammatory effects. In addition, the spatial arrangement of the electrical stimulation module enables localized electric field delivery, which promotes cell migration and proliferation, ultimately enhancing wound contraction. Meanwhile, the biosensing layer utilizes the upward-transported exudate to perform reliable monitoring of glucose, lactate, and pH under complex and dynamic wound conditions without interrupting the healing process. This vertically integrated modular design shows great potential for realizing miniaturized, high-density, scalable, and customizable multifunctional integration of therapy and monitoring. Various sensing and therapeutic units can be further integrated to meet the requirements of diverse clinical scenarios. Such versatility enables the platform to provide a feasible strategy for personalized wound care, improved therapeutic precision, and optimized clinical decision-making in the future applications.

## 4. Methods

### 4.1. Materials

All reagents and materials were used as received unless otherwise specified. Polyacrylonitrile (PAN), thermoplastic polyurethane (TPU), polyvinylpyrrolidone (PVP), pressure-sensitive adhesive (PSA), silver sulfadiazine, glucose oxidase (GOx), lactate oxidase (LOx), bovine serum albumin (BSA), tetrathiafulvalene (TTF), carbon nanotubes (CNTs), Nafion, chitosan, uric acid (UA), hydrochloric acid, anhydrous glucose, phosphate-buffered saline (PBS), polyaniline (PANI), and polyvinyl chloride (PVC) were purchased from Aladdin. Ultrapure water (18.2 MΩ·cm) was used for solution preparation. Dimethylformamide (DMF) was used as the solvent for electrospinning. PANI and PVC were obtained from Shanghai Acmec Biochemical Technology Co., Ltd. All other reagents were also sourced from Shanghai Acmec Biochemical Technology Co., Ltd.

### 4.2. HPAN layer

A 2 wt% PAN solution in DMF was prepared and electrospun under a voltage of 16 kV with the collector at –2 kV, a tip-to-collector distance of 13 cm, and a flow rate of 0.02 μL min⁻¹. The as-spun PAN membrane was hydrolyzed in a NaOH solution (3 g NaOH in 15 mL deionized water and 65 mL ethanol) at 50 °C for 15 min to obtain hydrolyzed PAN (HPAN). The HPAN membrane was dried under vacuum at 40 °C for 1 h prior to subsequent use.

### 4.3. TPU-PVP-PSA intermediate layer

A mixed solution containing 8.43 wt% TPU, 8.43 wt% PVP, and 33.3 wt% PSA in DMF was prepared and sonicated for 3 h to ensure homogeneity. Electrospinning was performed at 14 kV (collector at –2 kV), a tip-to-collector distance of 13 cm, and a flow rate of 0.035 μL min⁻¹.

### 4.4. TPU-PSA layer

A solution containing 9.0 wt% TPU and 50.0 wt% PSA in DMF was prepared and sonicated for 3 h. Electrospinning was conducted at 12.5 kV (collector at –2 kV), a tip-to-collector distance of 13 cm, and a flow rate of 0.04 μL min⁻¹.

### 4.5. Electrospinning environment

All electrospinning processes were performed at 50 °C and 40% relative humidity, using an electrospinning machine (TEADFS-100, Beijing Technova Technology Co., Ltd.)

### 4.6. Drug Loading

Silver sulfadiazine (AgSD) was incorporated into the TPU–PVP–PSA intermediate layer by adding it directly into the solution during preparation. Specifically, 0.5 g of AgSD was added for every 4.4 g of TPU and PVP. The mixture was stirred at 65 °C until it was fully homogenized.

### 4.7. Structural and Chemical Characterization

The surface and cross-sectional morphologies of the membranes were observed using scanning electron microscopy (SEM) (Zeiss EVO 18). Fiber diameters and pore sizes were measured from SEM images using ImageJ software (NIH). Fourier transform infrared (FTIR) spectra were recorded in transmission mode on a Nicolet iN10 instrument, with a scanning range from 4000 to 400 cm⁻¹. Water contact angles were determined using a goniometer (Biolin Scientific Theta Flex) by placing 3 μL droplets on the membrane surface. Mechanical properties, including tensile strength, were tested using an Instron universal testing machine (Instron 5566, Shimadzu EZ-LX) at a strain rate of 2 mm min⁻¹. Young’s modulus and strain at break were calculated from the stress–strain curves. The water evaporation rate was measured using a YG501B tester (Wenzhou Fangyuan Instrument Co., Ltd.) based on the GB/T12704.2-2009 standard. Moisture permeability tests were performed at 38 ± 2 °C and 50 ± 2 % relative humidity. The release of silver ions (Ag⁺) from the membrane was quantified using inductively coupled plasma-mass spectrometry (ICP-MS, iCAP RQ). Surface chemical composition and elemental states were analyzed by X-ray photoelectron spectroscopy (XPS, Kratos AXIS SUPRA+).

### 4.8. Fabrication of Patterned Electrodes

Aluminum plates were patterned using a UV picosecond laser cutter (LSP30) to create electrode masks. The electrodes for the electrostimulation and sensing modules were then fabricated via magnetron sputtering (SP-SC4-A00).

### 4.9. Fabrication of Sensor Arrays pH Sensor

The pH sensor was based on a polyaniline (PANI) film electrochemically polymerized onto a carbon nanotube (CNT) paper. The working electrode was cleaned by cyclic voltammetry (CV) in 0.5 M HCl with 10 cycles (scan rate: 0.1 V s⁻¹, potential range: −0.1 to 0.9 V). PANI electro-polymerization was performed in a 50 μL solution containing 0.1 M aniline and 1 M HCl using 50 CV cycles (−0.2 to 1.2 V, scan rate: 0.1 V s⁻¹). The CNT paper was then adhered to the gold (Au) electrode using conductive silver paste. The electrodes were air-dried at 4°C overnight.

## Glucose Sensor

A 0.1 M PBS solution was prepared by diluting 5 mL of 1× PBS with 45 mL deionized water. To fabricate the CNT electrode pad, 1 μL of a 0.1 M tetrathiafulvalene (TTF) solution (0.02 g TTF dissolved in 1 mL anhydrous ethanol) was drop-cast onto the pad. The enzyme coating solution was prepared by dissolving 40 mg/mL glucose oxidase (GOx) in 0.1 M PBS containing 10 mg/mL bovine serum albumin (BSA), with a final GOx concentration of 100 mg/mL. A 1 wt% Nafion solution was prepared by mixing 1 mL of 5% Nafion with 1 mL anhydrous ethanol, followed by dilution to 2 mL with 0.1 M PBS. The Nafion solution was mixed in equal volume with the enzyme solution and drop-cast onto the TTF-coated CNT pad. The CNT pad was then adhered to a gold current collector using conductive silver paste, and the assembly was cured at 4°C for 24 hours. The electrodes were stored at 4°C for at least 1 week to allow the Nafion membrane to equilibrate.

## Lactate Sensor

The lactate sensor was fabricated following a similar procedure to the glucose sensor. In addition to the glucose oxidase (GOx) enzyme layer, a 3 μL chitosan solution (prepared by dissolving 1% chitosan in 2% acetic acid and stirring for 1 hour) was drop-cast onto the electrode surface. After the chitosan layer, 3 μL of polyvinyl chloride (PVC) solution was applied on top of the chitosan layer.

## Reference Electrode

The Ag/AgCl reference electrode was prepared by directly applying Ag/AgCl gel onto a gold electrode and allowing it to air dry.

## Counter Electrode

For the counter electrode, a 10% platinum-carbon (Pt/C) suspension (10 mg/mL) was prepared in a 2 wt% Nafion ethanol solution. 5 μL of the Pt/C suspension was drop-cast onto the current collector.

### 4.10. In vitro cell studies Cell Lines

NIH-3T3 fibroblasts were cultured under 37°C and 5% CO₂. Cells were passaged at 70% confluency, with passages 3 to 5 used for all studies.

## In vitro biocompatibility and Proliferation Studies

Electroactive hydrogels were washed and transferred to 24-well cell culture inserts. NIH-3T3 fibroblasts were seeded at a density of 1 × 10⁵ cells per well. The inserts were placed in 24-well plates, and cells were incubated with appropriate media at 37°C and 5% CO₂. Cell proliferation and viability were assessed using a commercial PrestoBlue assay (Thermo Fisher Scientific) to evaluate metabolic activity. For cell viability, a calcein AM/ethidium homodimer-1 live/dead kit (Invitrogen) was used. Live cells were stained green with calcein-AM, and dead cells were stained red with ethidium homodimer-1. The percentage of live cells was calculated using ImageJ software by determining the ratio of live cells to total cells.

### 4.11. In Vivo Wound Healing Model Animal Model

In vivo studies were conducted on Sprague-Dawley rats (female) to assess the wound healing efficacy of the trilayer dressing. The rats were anesthetized with isoflurane (2-3% in oxygen), and full-thickness dorsal skin defects (0.8 cm diameter) were induced on their backs.

## Electrical stimulation therapy

The stimulation protocol was preprogrammed and kept identical across animals, consisting of a 10 min stimulation phase followed by a 20 min rest period. During each stimulation phase, a 1 V, 50 Hz signal was applied to deliver consistent low-voltage electrical stimulation at the wound interface.

## Wound Closure and Epithelialization

The wound closure rate was quantified by analyzing the wound area using image processing. Epithelialization and epidermal thickness were evaluated using hematoxylin and eosin (HE) staining. The epidermal thickness was measured on histological sections from the wound area.

## Histological Staining and Analysis

To assess tissue remodeling and myofibroblast activation, histological staining for α-SMA and Masson trichrome was performed on tissue sections. Sections were incubated with primary antibodies specific for α-SMA, and Masson trichrome was used to stain collagen. The expression levels of α-SMA and the percentage of collagen-positive areas were quantified using ImageJ. The areas of positive staining were calculated to determine myofibroblast presence and collagen content, providing insights into tissue regeneration and inflammation during the wound healing process.

## Animal Infection Model

5 μL of Staphylococcus aureus (S. aureus) bacterial solution was applied to the wound site on the dorsal area of the mice five times. After each application, the solution was allowed to air dry before the next application.

## Biochemical Parameter Monitoring

Wound fluid samples were collected for continuous biochemical analysis of glucose, lactate, and pH. The concentrations were measured using biosensors integrated into the dressing. Glucose and lactate were monitored using specific sensors, and pH was measured using a pH sensor. Wound fluid was collected daily for 5 days, and parameters were recorded at regular intervals to track the biochemical changes during healing.

## Data availability

All data needed to support the conclusions of this manuscript are included in the main text and the supplementary information/source data file and are available from the corresponding authors upon request.

## References

(1) Queen, D.; Harding, K. What’s the true costs of wounds faced by different healthcare systems around the world? Int Wound J 2023, 20 (10), 3935–3938.

(2) Rodrigues, M.; Kosaric, N.; Bonham, C. A.; Gurtner, G. C. Wound Healing: A Cellular Perspective. Physiol Rev 2019, 99 (1), 665–706.

(3) Tracy, L. E.; Minasian, R. A.; Caterson, E. J. Extracellular Matrix and Dermal Fibroblast Function in the Healing Wound. Adv Wound Care 2016, 5 (3), 119–136.

(4) Eming, S. A.; Martin, P.; Tomic-Canic, M. Wound repair and regeneration: Mechanisms, signaling, and translation. Sci Transl Med 2014, 6 (265).

(5) Armstrong, D. G.; Boulton, A. J. M.; Bus, S. A. Diabetic Foot Ulcers and Their Recurrence. New Engl J Med 2017, 376 (24), 2367–2375.

(6) Shaydakov, M. E.; Ting, W.; Sadek, M.; Aziz, F.; Diaz, J. A.; Raffetto, J. D.; Marston, W. A.; Lal, B. K.; Welch, H. J.; Comm, A. V. F. R. Review of the current evidence for topical treatment for venous leg ulcers. J Vasc Surg-Venous L 2022, 10 (1), 241–247.

(7) Saramago, P.; Gkekas, A.; Arundel, C. E.; Chetter, I. C.; Investigators, S.-T. Negative pressure wound therapy for surgical wounds healing by secondary intention is not cost-effective. Bjs-Brit J Surg 2025, 112 (5).

(8) Aslam, B.; Wang, W.; Arshad, M. I.; Khurshid, M.; Muzammil, S.; Rasool, M. H.; Nisar, M. A.; Alvi, R. F.; Aslam, M. A.; Qamar, M. U.;, et al. Antibiotic resistance: a rundown of a global crisis. Infect Drug Resist 2018, 11, 1645–1658.

(9) Dong, R. N.; Guo, B. L. Smart wound dressings for wound healing. Nano Today 2021, 41.

(10) Farahani, M.; Shafiee, A. Wound Healing: From Passive to Smart Dressings. Adv Healthc Mater 2021, 10 (16).

(11) Zhang, W.; Liu, L. L.; Cheng, H.; Zhu, J.; Li, X. Y.; Ye, S.; Li, X. J. Hydrogel-based dressings designed to facilitate wound healing. Mater Adv 2024, 5 (4), 1364–1394.

(12) Mostafalu, P.; Tamayol, A.; Rahimi, R.; Ochoa, M.; Khalilpour, A.; Kiaee, G.; Yazdi, I. K.; Bagherifard, S.; Dokmeci, M. R.; Ziaie, B.;, et al. Smart Bandage for Monitoring and Treatment of Chronic Wounds. Small 2018, 14 (33).

(13) Wang, M. Q.; Yang, Y. R.; Min, J. H.; Song, Y.; Tu, J. B.; Mukasa, D.; Ye, C.; Xu, C. H.; Heflin, N.; McCune, J. S.;, et al. A wearable electrochemical biosensor for the monitoring of metabolites and nutrients. *Nat*. Biomed. Eng. 2022, 6 (11), 1225–1235.

(14) Sani, E. S.; Xu, C. H.; Wang, C. R.; Song, Y.; Min, J. H.; Tu, J. B.; Solomon, S. A.; Li, J. H.; Banks, J. L.; Armstrong, D. G.;, et al. A stretchable wireless wearable bioelectronic system for multiplexed monitoring and combination treatment of infected chronic wounds. Sci Adv 2023, 9 (12).

(15) Zhang, X.; Lv, R.; Chen, L.; Sun, R.; Zhang, Y.; Sheng, R.; Du, T.; Li, Y.; Qi, Y. A Multifunctional Janus Electrospun Nanofiber Dressing with Biofluid Draining, Monitoring, and Antibacterial Properties for Wound Healing. ACS Appl Mater Interfaces 2022, 14 (11), 12984–13000.

(16) Wang, C. R.; Fan, K. X.; Sani, E. S.; Lasalde-Ramírez, J. A.; Heng, W. Z.; Min, J. H.; Solomon, S. A.; Wang, M. Q.; Li, J. H.; Han, H.;, et al. A microfluidic wearable device for wound exudate management and analysis in human chronic wounds. Sci Transl Med 2025, 17 (795).

(17) Shi, C. X.; Wang, H.; Wang, X. J.; Li, K. F.; Liu, P. B.; Wang, L. H.; Yu, H. A Safe, Stable, Simple, Serviceable, and Self-Powered Wound Dressing With Continuous Low-Voltage Direct Current Electrical Stimulation: an Efficient Approach to Accelerate Wound Healing. Adv Funct Mater 2025, 35 (29).

(18) Peng, Q. Y.; Qian, Y.; Xiao, X. X.; Gao, F. D.; Ren, G. Q.; Pennisi, C. P. Advancing Chronic Wound Healing through Electrical Stimulation and Adipose-Derived Stem Cells. Adv Healthc Mater 2025, 14 (10).

(19) Zhao, M.; Song, B.; Pu, J.; Wada, T.; Reid, B.; Tai, G. P.; Wang, F.; Guo, A. H.; Walczysko, P.; Gu, Y.;, et al. Electrical signals control wound healing through phosphatidylinositol-3-OH kinase-γ and PTEN. Nature 2006, 442 (7101), 457–460.

(20) Liu, Y. R.; Huang, S. F.; Liang, S. Y.; Lin, P. R.; Lai, X. J.; Lan, X. Z.; Wang, H.; Tang, Y. D.; Gao, B. T. Phase Change Material-Embedded Multifunctional Janus Nanofiber Dressing with Directional Moisture Transport, Controlled Release of Anti-Inflammatory Drugs, and Synergistic Antibacterial Properties. Acs Appl Mater Inter 2023, 15 (45), 52244–52261.

(21) Wang, S.; Wang, N. N.; Kai, D.; Li, B. F.; Wu, J.; Yeo, J. C. C.; Xu, X. W.; Zhu, J.; Loh, X. J.; Hadjichristidis, N.;, et al. In-situ forming dynamic covalently crosslinked nanofibers with one-pot closed-loop recyclability. Nat Commun 2023, 14 (1).

(22) Chen, B.; He, B.; Tucker, A. M.; Biluck, I.; Leung, T. H.; Schaer, T. P.; Yang, S. An Environmentally Stable, Biocompatible, and Multilayered Wound Dressing Film with Reversible and Strong Adhesion. Adv Healthc Mater 2024, 13 (31).

(23) Xu, G.; Lu, Y. L.; Cheng, C.; Li, X.; Xu, J.; Liu, Z. Y.; Liu, J. L.; Liu, G.; Shi, Z. H.; Chen, Z. T.;, et al. Battery-Free and Wireless Smart Wound Dressing for Wound Infection Monitoring and Electrically Controlled On-Demand Drug Delivery. Adv Funct Mater 2021, 31 (26).

(24) Ge, Z. Y.; Guo, W. S.; Tao, Y.; Sun, H. X.; Meng, X. Y.; Cao, L. Y.; Zhang, S. G.; Liu, W. Y.; Akhtar, M. L.; Li, Y.;, et al. Wireless and Closed-Loop Smart Dressing for Exudate Management and On-Demand Treatment of Chronic Wounds. Adv Mater 2023, 35 (47).

(25) Deng, W.; Sun, M. M.; Cao, M. Z.; Ma, C. B.; Bo, X. J.; Bai, J.; Zhou, M. A Fully Integrated Wearable Biomimetic Microfluidic Wound Tracker for In Situ Dynamic Monitoring of Wound Exudate Oxygen. Acs Nano 2025, 19 (16), 16163–16174.

(26) Zhang, X. M.; Lv, R. J.; Chen, L. X.; Sun, R. M.; Zhang, Y.; Sheng, R. T.; Du, T.; Li, Y. H.; Qi, Y. F. A Multifunctional Janus Electrospun Nanofiber Dressing with Biofluid Draining, Monitoring, and Antibacterial Properties for Wound Healing. Acs Appl Mater Inter 2022, 14 (11), 12984–13000.

(27) Wang, H. B.; Wang, M. J.; Xu, X. H.; Gao, P.; Xu, Z. L.; Zhang, Q.; Li, H. Y.; Yan, A. X.; Kao, R. Y. T.; Sun, H. Z. Multi-target mode of action of silver against Staphylococcus aureus endows it with capability to combat antibiotic resistance. Nat Commun 2021, 12 (1).

(28) Dong, Y. P.; Fu, S. J.; Yu, J. Y.; Li, X. R.; Ding, B. Emerging Smart Micro/Nanofiber-Based Materials for Next-Generation Wound Dressings. Adv Funct Mater 2024, 34 (9).

(29) Qian, L. L.; Jin, F.; Li, T.; Wei, Z. D.; Ma, X. Y.; Zheng, W. Y.; Javanmardi, N.; Wang, Z.; Ma, J.; Lai, C. T.;, et al. Self-Adhesive and Self-Sustainable Bioelectronic Patch for Physiological Feedback Electronic Modulation of Soft Organs. Adv Mater 2024, 36 (41).

(30) Qian, S. T.; Wang, J.; Liu, Z. M.; Mao, J. Y.; Zhao, B. F.; Mao, X. Y.; Zhang, L. C.; Cheng, L. Y.; Zhang, Y. G.; Sun, X. M.;, et al. Secretory Fluid-Aggregated Janus Electrospun Short Fiber Scaffold for Wound Healing. Small 2022, 18 (36).

(31) Shi, C. X.; Wang, H.; Wang, X. J.; Li, K. F.; Liu, P. B.; Wang, L. H.; Yu, H. A Safe, Stable, Simple, Serviceable, and Self-Powered Wound Dressing With Continuous Low-Voltage Direct Current Electrical Stimulation: an Efficient Approach to Accelerate Wound Healing. Adv Funct Mater 2025.

(32) Zhang, S. C.; Liu, H.; Tang, N.; Ge, J. L.; Yu, J. Y.; Ding, B. Direct electronetting of high-performance membranes based on self-assembled 2D nanoarchitectured networks. Nat Commun 2019, 10.

(33) Ballengee, J. B.; Pintauro, P. N. Morphological Control of Electrospun Nafion Nanofiber Mats. J Electrochem Soc 2011, 158 (5), B568–B572.

(34) Ji, D. X.; Lin, Y. G.; Guo, X. Y.; Ramasubramanian, B.; Wang, R. W.; Radacsi, N.; Jose, R.; Qin, X. H.; Ramakrishna, S. Electrospinning of nanofibres. *Nat Rev Method Prime* 2024, 4 (1).

(35) Keirouz, A.; Wang, Z.; Reddy, V. S.; Nagy, Z. K.; Vass, P.; Buzgo, M.; Ramakrishna, S.; Radacsi, N. The History of Electrospinning: Past, Present, and Future Developments. Adv Mater Technol-Us 2023, 8 (11).

(36) Wu, L. H.; Wang, Y. Q.; Zhao, X. Y.; Zhao, T. T.; Li, J. H.; Kuang, Y.; He, Y. Y.; Yang, S. X.; Gu, Z. W.; Mao, H. L. A self-adhesive hierarchical nanofiber patch for dynamic and multistage management of full-thickness cutaneous wounds. J Nanobiotechnol 2025, 23 (1).

(37) Yu, H.; Li, Y. J.; Pan, Y. N.; Wang, H. N.; Wang, W.; Ren, X. B.; Yuan, H.; Lv, Z. R.; Zuo, Y. J.; Liu, Z. R.;, et al. Multifunctional porous poly (L-lactic acid) nanofiber membranes with enhanced anti-inflammation, angiogenesis and antibacterial properties for diabetic wound healing. J Nanobiotechnol 2023, 21 (1).

(38) Sun, L. L.; Wang, J. C.; Matsui, H.; Lee, S. Y.; Wang, W. Q.; Guo, S. Y.; Chen, H. T.; Fang, K.; Ito, Y.; Inoue, D.;, et al. All-solution-processed ultraflexible wearable sensor enabled with universal trilayer structure for organic optoelectronic devices. Sci Adv 2024, 10 (15).

(39) Kang, T. W.; Lee, Y. J.; Rigo, B.; Soltis, I.; Lee, J.; Kim, H.; Wang, G. R.; Zavanelli, N.; Ayesh, E.; Sohail, W.;, et al. Soft Nanomembrane Sensor-Enabled Wearable Multimodal Sensing and Feedback System for Upper-Limb Sensory Impairment Assistance. Acs Nano 2025, 19 (5), 5613–5628.

(40) He, X. C.; Wang, W. Y.; Yang, S. J.; Zhang, F. L.; Gu, Z.; Dai, B.; Xu, T. L.; Huang, Y. Y. S.; Zhang, X. J. Adhesive tapes: From daily necessities to flexible smart electronics. Appl Phys Rev 2023, 10 (1).

(41) Creton, C. Pressure-sensitive adhesives: An introductory course. MRS Bull. 2003, 28 (6), 434–439.

(42) Fitzgerald, D. M.; Colson, Y. L.; Grinstaff, M. W. Synthetic pressure sensitive adhesives for biomedical applications. Prog Polym Sci 2023, 142.

(43) Yang, X.; Liu, X. N.; Chau, Y. Y.; Qin, X. Z.; Zhu, H.; Peng, L.; Chan, K. W. Y.; Wang, Z. K. Role of chemistry in nature-inspired skin adhesives. Chem Sci 2025, 16 (24), 10665–10690.

(44) Kudryavtsev, Y. V.; Gelinck, E.; Fischer, H. R. Theoretical investigation of van der Waals forces between solid surfaces at nanoscales. Surf Sci 2009, 603 (16), 2580–2587.

(45) Ciavarella, M.; Joe, J.; Papangelo, A.; Barber, J. R. The role of adhesion in contact mechanics. J R Soc Interface 2019, 16 (151).

(46) Geng, C.; He, S.; Yu, S.; Johnson, H. M.; Shi, H. X.; Chen, Y. B.; Chan, Y. K.; He, W. X.; Qin, M.; Li, X.;, et al. Achieving Clearance of Drug-Resistant Bacterial Infection and Rapid Cutaneous Wound Regeneration Using an ROS-Balancing-Engineered Heterojunction. Adv. Mater. 2024.

(47) Wu, L.; Huang, X.; Wang, M.; Chen, J. S. Z.; Chang, J. K.; Zhang, H.; Zhang, X. T.; Conn, A.; Rossiter, J.; Birchall, M.;, et al. Tunable Light-Responsive Polyurethane-urea Elastomer Driven by Photochemical and Photothermal Coupling Mechanism. Acs Appl Mater Inter 2024, 16 (15), 19480–19495.

(48) Paez-Amieva, Y.; Mateo-Oliveras, N.; Martín-Martínez, J. M. Polyurethanes Synthesized with Blends of Polyester and Polycarbonate Polyols-New Evidence Supporting the Dynamic Non-Covalent Exchange Mechanism of Intrinsic Self-Healing at 20 °C. Polymers-Basel 2024, 16 (20).

(49) Laukkanen, T.; Reddy, P. G.; Barua, A.; Kumar, M.; Kolpakov, K.; Tirri, T.; Sharma, V. Sustainable castor oil-derived cross-linked poly(ester-urethane) elastomeric films for stretchable transparent conductive electrodes and heaters. J Mater Chem A 2024, 12 (47), 33177–33192.

(50) Wang, H. R.; Cao, L.; Wang, X. L.; Lang, X. R.; Cong, W. W.; Han, L.; Zhang, H. Y.; Zhou, H. B.; Sun, J. J.; Zong, C. Z. Effects of Isocyanate Structure on the Properties of Polyurethane: Synthesis, Performance, and Self-Healing Characteristics. Polymers-Basel 2024, 16 (21).

(51) Wong, S. H. D.; Deen, G. R.; Bates, J. S.; Maiti, C.; Lam, C. Y. K.; Pachauri, A.; AlAnsari, R.; Belsky, P.; Yoon, J.; Dodda, J. M. Smart Skin-Adhesive Patches: From Design to Biomedical Applications. Adv Funct Mater 2023, 33 (14).

(52) Yang, Z. X.; Ren, K. X.; Chen, Y. H.; Quanji, X. Y.; Cai, C. F.; Yin, J. B. Oxygen-Generating Hydrogels as Oxygenation Therapy for Accelerated Chronic Wound Healing. Adv Healthc Mater 2024, 13 (3).

(53) Wang, X. F.; Huang, Z.; Miao, D. Y.; Zhao, J.; Yu, J. Y.; Ding, B. Biomimetic Fibrous Murray Membranes with Ultrafast Water Transport and Evaporation for Smart Moisture-Wicking Fabrics. Acs Nano 2019, 13 (2), 1060–1070.

(54) Kleinbeck, K. R.; Bader, R. A.; Kao, W. J. Concurrent In Vitro Release of Silver Sulfadiazine and Bupivacaine From Semi-Interpenetrating Networks for Wound Management. J Burn Care Res 2009, 30 (1), 98–104.

(55) Bergonzi, C.; Bianchera, A.; Remaggi, G.; Ossiprandi, M. C.; Bettini, R.; Elviri, L. 3D Printed Chitosan/Alginate Hydrogels for the Controlled Release of Silver Sulfadiazine in Wound Healing Applications: Design, Characterization and Antimicrobial Activity. Micromachines-Basel 2023, 14 (1).

(56) Jung, W. K.; Koo, H. C.; Kim, K. W.; Shin, S.; Kim, S. H.; Park, Y. H. Antibacterial activity and mechanism of action of the silver ion in Staphylococcus aureus and Escherichia coli. Appl Environ Microb 2008, 74 (7), 2171–2178.

(57) Liao, C. Z.; Li, Y. C.; Tjong, S. C. Bactericidal and Cytotoxic Properties of Silver Nanoparticles. Int J Mol Sci 2019, 20 (2).

(58) Zhu, Y. N.; Zhang, J. M.; Song, J. Y.; Yang, J.; Du, Z.; Zhao, W. Q.; Guo, H. S.; Wen, C. Y.; Li, Q. S.; Sui, X. J.;, et al. A Multifunctional Pro-Healing Zwitterionic Hydrogel for Simultaneous Optical Monitoring of pH and Glucose in Diabetic Wound Treatment. Adv Funct Mater 2020, 30 (6).

(59) Huang, X. C.; Zhang, J. R.; Zhang, L. L.; Su, H.; Liu, X. M.; Liu, J. A sensitive H2O2 biosensor based on carbon nanotubes/tetrathiafulvalene and its application in detecting NADH. Anal Biochem 2020, 589.

(60) Mecheri, B.; De Porcellinis, D.; Campana, P. T.; Rainer, A.; Trombetta, M.; Marletta, A.; Oliveira, O. N.; Licoccia, S. Tuning Structural Changes in Glucose Oxidase for Enzyme Fuel Cell Applications. Acs Appl Mater Inter 2015, 7 (51), 28311–28318.

(61) Garcia-Guzmán, J. J.; Heras, J. M. J.; López-Iglesias, D.; Gonzalez-alvarez, R. J.; Cubillana-Aguilera, L.; Macias, C. G.; Alba, J. J. F.; Palacios-Santander, J. M. New spin coated multilayer lactate biosensor for acidosis monitoring in continuous flow assisted with an electrochemical pH probe. Microchim Acta 2024, 191 (9).

(62) Lyu, Y.; Mollik, P.; Olah, A. L.; Halter, D. P. Construction and Evaluation of Cheap and Robust Miniature Ag/AgCl Reference Electrodes for Aqueous and Organic Electrolytes. Chemelectrochem 2024, 11 (8).

(63) Leal, J.; Shaner, S.; Jedrusik, N.; Savelyeva, A.; Asplund, M. Electrotaxis evokes directional separation of co-cultured keratinocytes and fibroblasts. Sci Rep-Uk 2023, 13 (1).

(64) Song, B.; Zhao, M.; Forrester, J. V.; McCaig, C. D. Electrical cues regulate the orientation and frequency of cell division and the rate of wound healing. Proceedings of the National Academy of Sciences of the United States of America 2002, 99 (21), 13577–13582.

(65) Nakajima, K.; Zhu, K.; Sun, Y. H.; Hegyi, B.; Zeng, Q. L.; Murphy, C. J.; Small, J. V.; Ye, C. I.; Izumiya, Y.; Penninger, J. M.;, et al. /Kir4.2 couples with polyamines to sense weak extracellular electric fields in galvanotaxis. Nat. Commun. 2015, 6.

(66) Lin, B. J.; Tsao, S. H.; Chen, A.; Hu, S. K.; Chao, L.; Chao, P. H. G. Lipid rafts sense and direct electric field-induced migration. P Natl Acad Sci USA 2017, 114 (32), 8568–8573.

(67) Maslova, E.; EisaianKhongi, L.; Rigole, P.; Coenye, T.; McCarthy, R. R. Carbon source competition within the wound microenvironment can significantly influence infection progression. Npj Biofilms Microbi 2024, 10 (1).

(68) Khatib-Massalha, E.; Bhattacharya, S.; Massalha, H.; Biram, A.; Golan, K.; Kollet, O.; Kumari, A.; Avemaria, F.; Petrovich-Kopitman, E.; Gur-Cohen, S.;, et al. Lactate released by inflammatory bone marrow neutrophils induces their mobilization via endothelial GPR81 signaling. Nat Commun 2020, 11 (1).

(69) Zhao, X.; Huang, J. H.; Zhang, J. C.; Yang, B. W.; Hu, Z. J.; Li, T.; Ma, X.; Jiang, C. Y.; Zou, H. C.; Liu, S. R.;, et al. Soft bioelectronics embedded with self-confined tetrahedral DNA circuit for high-fidelity chronic wound monitoring. Nat Commun 2025, 16 (1).

(70) Haller, H. L.; Sander, F.; Popp, D.; Rapp, M.; Hartmann, B.; Demircan, M.; Nischwitz, S. P.; Kamolz, L. P. Oxygen, pH, Lactate, and Metabolism-How Old Knowledge and New Insights Might Be Combined for New Wound Treatment. Medicina-Lithuania 2021, 57 (11).

(71) Dunker, C.; Polke, M.; Schulze-Richter, B.; Schubert, K.; Rudolphi, S.; Gressler, A. E.; Pawlik, T.; Prada Salcedo, J. P.; Niemiec, M. J.; Slesiona-Künzel, S.;, et al. Rapid proliferation due to better metabolic adaptation results in full virulence of a filament-deficient Candida albicans strain. Nat Commun 2021, 12 (1).

