## Supplementary Information for "Exudate-Guided Janus Trilayer Bioelectronic Dressing for Multiplexed Sensing and Therapy of Chronic Wounds"

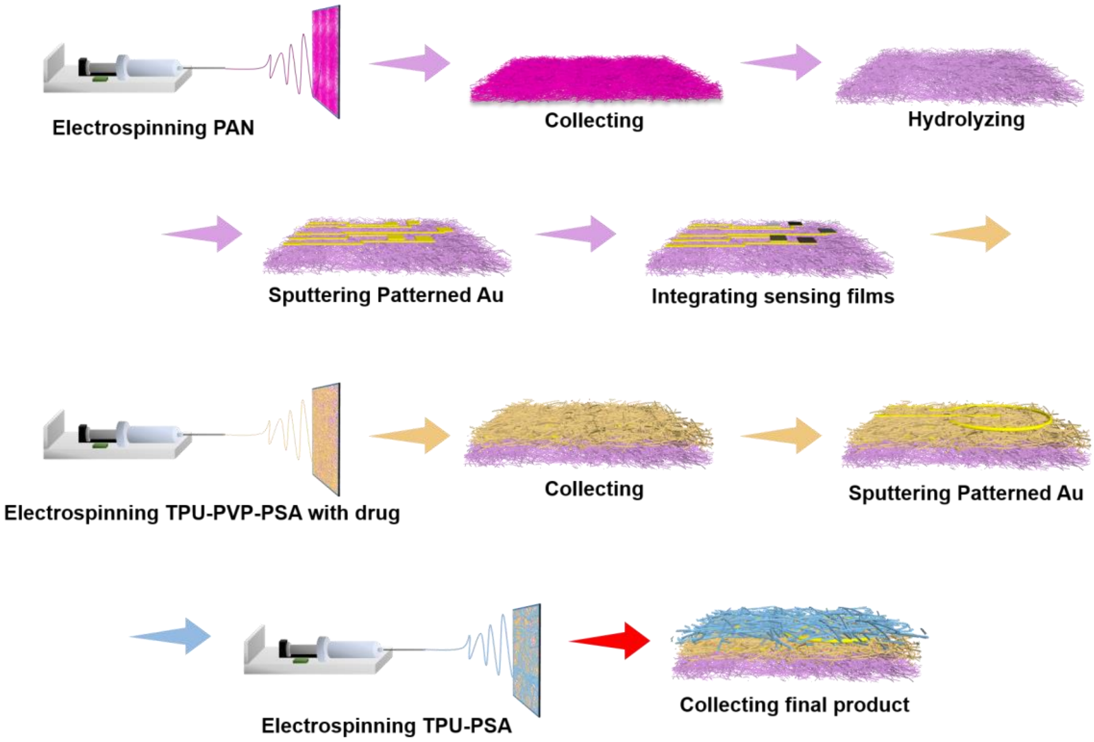


**Figure S1.** The fabrication process of the bioelectronic wound dressing.

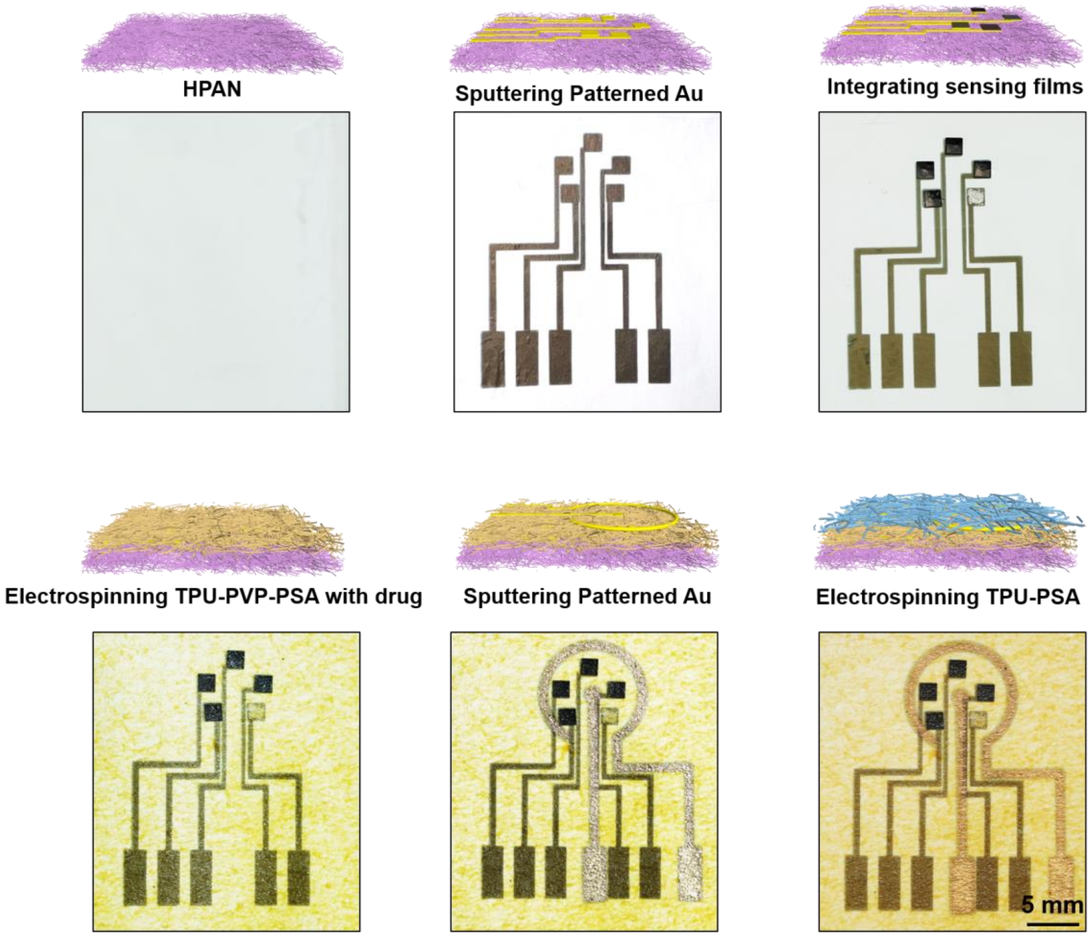


**Figure S2.** Photographic images of each fabrication step in the preparation of the trilayer bioelectronic wound dressing.

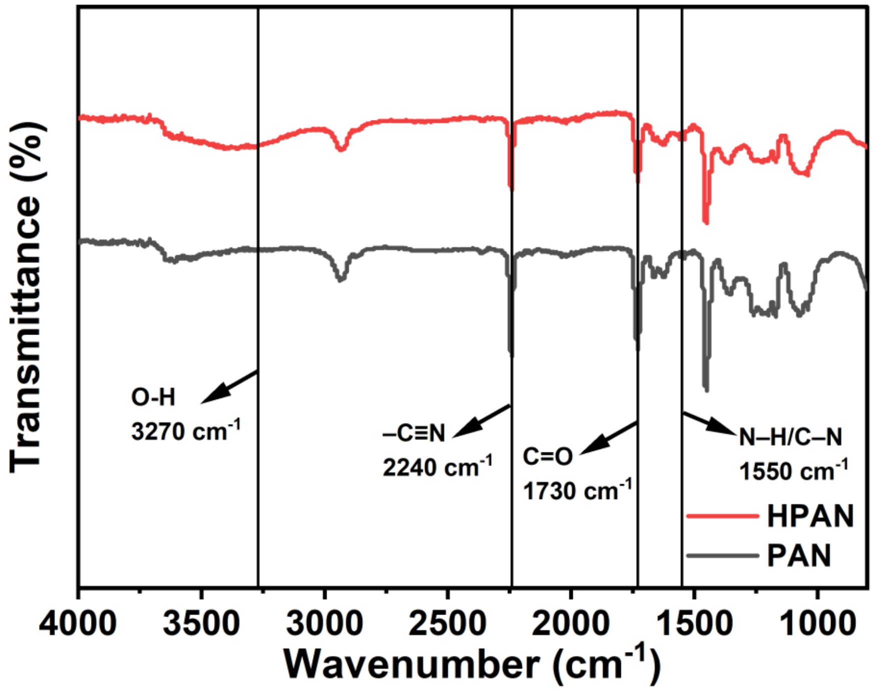


**Figure S3.** FT-IR spectra of pristine PAN and hydrolyzed HPAN.

Key spectral transitions confirm successful hydrolysis/ Emergence of broad O-H stretch at 3270 cm$^{-1}$, enhanced C=O stretch at 1730 cm$^{-1}$ (carboxylic acid formation) and Characteristic amide II band at 1550 cm$^{-1}$ (coupled N-H bending and C-N stretching), and concomitant reduction in -C≡N peak intensity at 2240 cm$^{-1}$ (nitrile group consumption). This collective shift verifies the generation of hydrophilic hydroxyl/carboxyl functionalities at the expense of hydrophobic nitrile groups, directly correlating with the significantly enhanced hydrophilicity of HPAN.

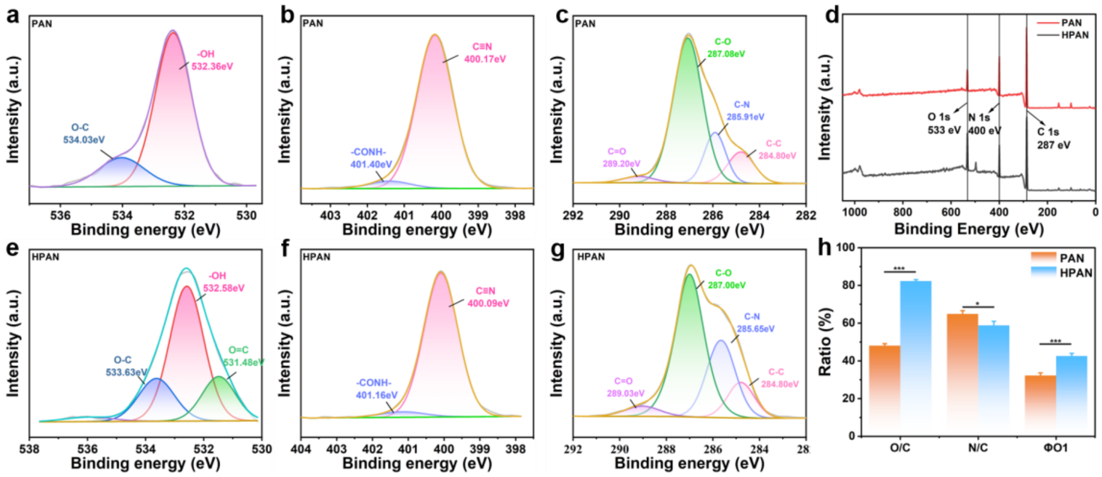


**Figure S4.** XPS analysis of PAN and hydrolyzed HPAN. (a-c) High-resolution XPS spectra of pristine PAN showing the (a) O 1s, (b) N 1s, and (c) C 1s core levels. (d) Survey scan spectra comparing the elemental composition of pristine PAN and HPAN. (e-g) High-resolution XPS spectra of HPAN showing the (e) O 1s, (f) N 1s, and (g) C 1s core levels. (h) Quantitative analysis of elemental composition ratios (O/C, N/C) and O 1s chemical state changes (φO1, representing the fraction of C=O plus C–O/–OH species relative to total O 1s peak area) induced by hydrolysis.

Core-level XPS spectra document the hydrolysis-induced chemical transformation: The O 1s spectrum of pristine PAN (Figure S4a) exhibits characteristic peaks at 532.36 eV (hydroxyl groups) and 534.03 eV (ether-type oxygen), while its N 1s spectrum (Figure S4b) shows signatures at 400.17 eV (nitrile nitrogen) and 401.40 eV (amide nitrogen). Corresponding C 1s signals (Figure S4c) appear at 284.80 eV (aliphatic carbon), 285.91 eV (carbon-nitrogen bonds), 287.08 eV (alcohol/ether carbon), and 289.20 eV (carbonyl carbon). Following hydrolysis, HPAN demonstrates significant chemical shifts: The O 1s spectrum (Figure S4e) displays new components at 531.48 eV (carboxylic oxygen) alongside modified peaks at 532.58 eV (hydroxyl) and 534.63 eV (ether oxygen); the N 1s spectrum (Figure S4f) shows attenuated nitrile intensity at 400.10 eV with persistent amide contribution at 401.16 eV; the C 1s spectrum (Figure S4g) maintains the carbon skeleton signal at 284.80 eV but reveals chemical environment alterations at 285.65 eV (C-N), 287.00 eV (C-O), and 289.03 eV (C=O). Full-survey spectra (Figure S4d) confirm elemental redistribution: nitrogen signal attenuation directly attributable to nitrile group consumption contrasts with oxygen signal enhancement from newly generated hydroxyl and carboxyl functionalities, while carbon peak broadening indicates heteroatom incorporation into the polymer backbone. Quantitative analysis (Figure S4h) reveals substantially increased oxygen-to-carbon atomic ratio and elevated proportion of polar carbonyl/hydroxyl groups within O 1s spectra, coupled with decreased nitrogen-to-carbon ratio, collectively evidencing successful conversion of hydrophobic nitriles to hydrophilic moieties that significantly enhance surface wettability.

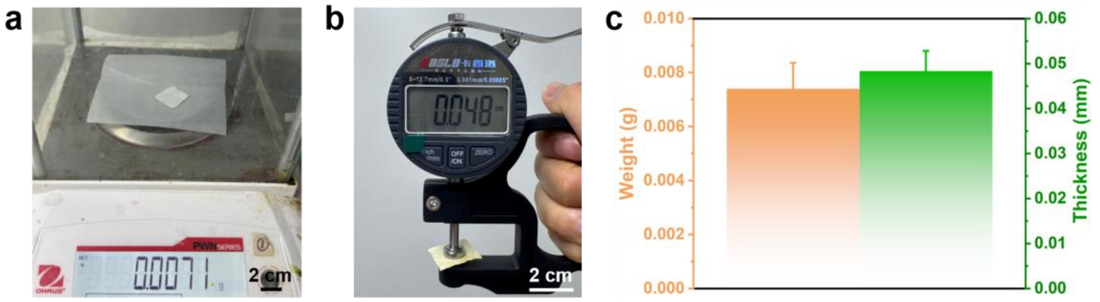


**Figure S5.** Photograph images of the trilayer (a) with a total weight of 0.0071 g (b) and thickness of 0.048 mm. (c) Average weight and thickness of trilayer. The dimensions of the samples were fixed as 2 cm by 2 cm. Error bars represent mean ± SD (n=3).

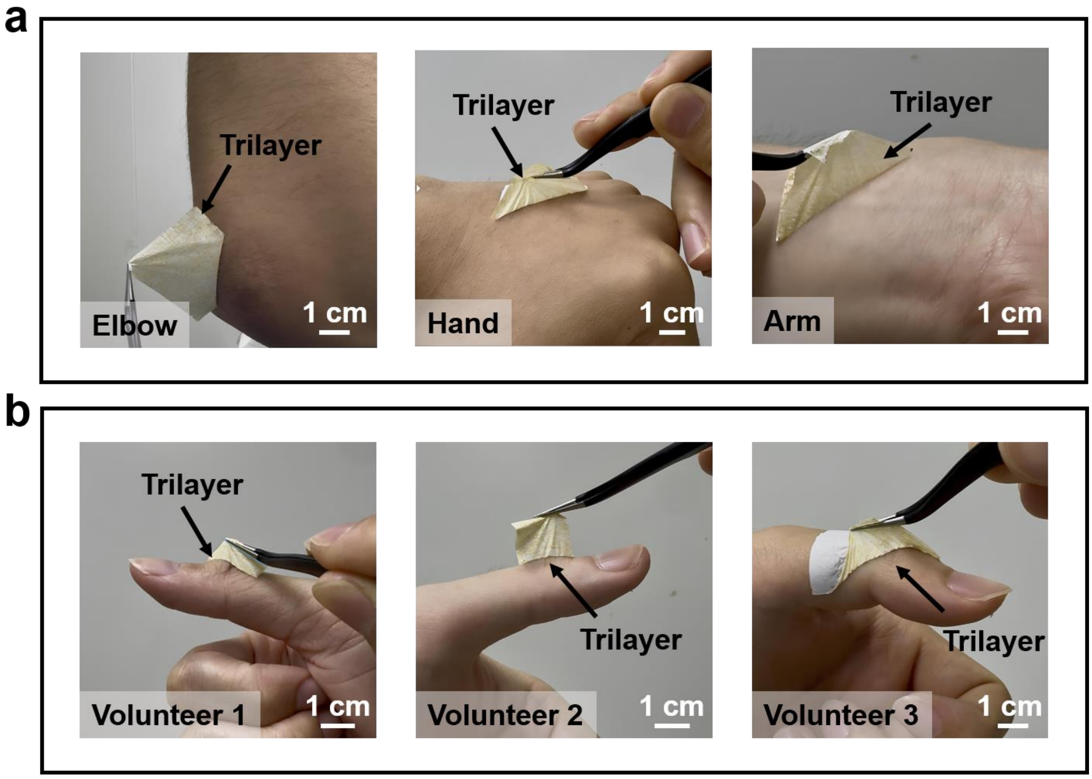


**Figure S6.** Surface adhesion properties of the trilayer membrane. Photos showing the trilayer membrane detached from (a) different parts of the body (b) and the finger joints of different volunteers without any harms.

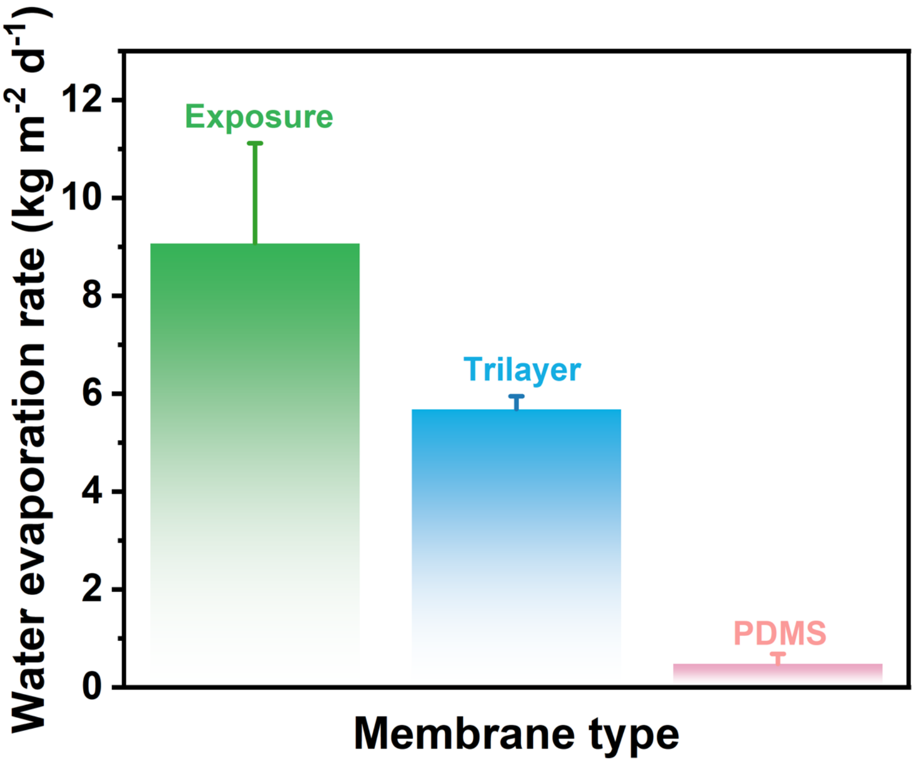


**Figure S7.** Statistical results of quantitative data on the breathability of the trilayer membrane. Error bars represent mean ± SD (n=3).

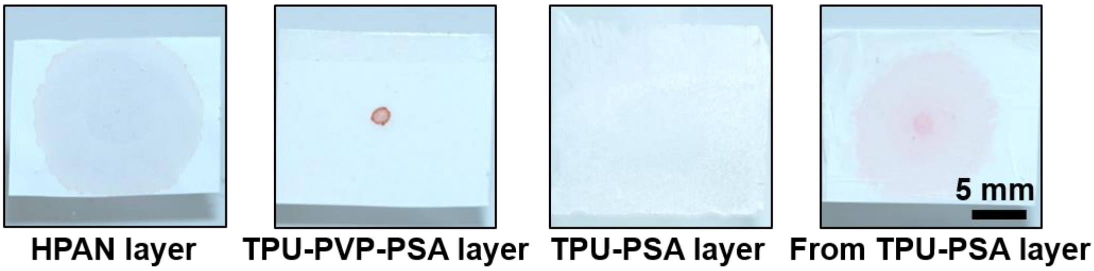


**Figure S8.** Top-view optical images depicting the spreading behavior of 3 μL water droplets after 5 minutes on each individual layer and the trilayer membrane.

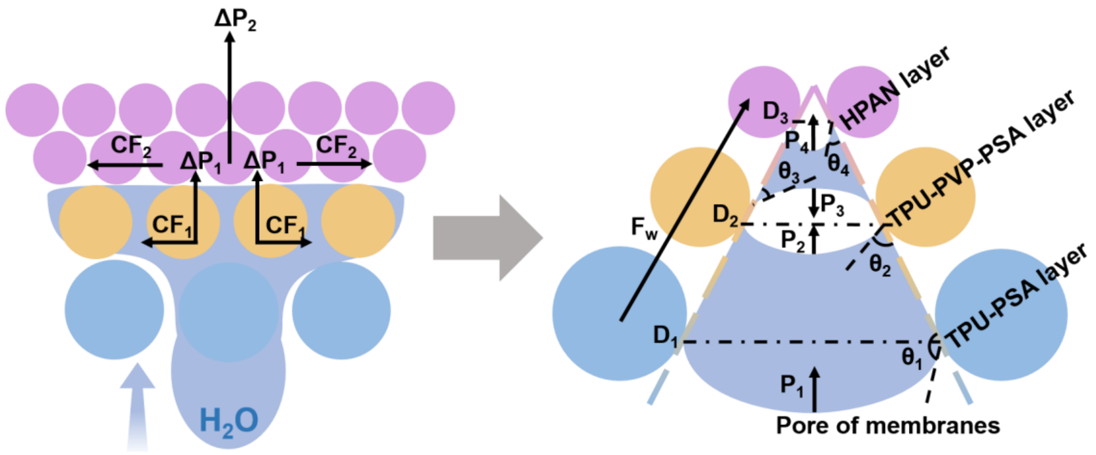


**Figure S9.** Diagram of the simplified mechanism of passive wicking process of trilayer membrane.

When a water droplet initially contacts the trilayer membrane, it rapidly infiltrates into the inter-fiber capillary channels of the hydrophobic TPU-PSA layer due to localized wetting at fiber junctions. Subsequently, the liquid is subjected to Laplace pressure, described as:

$$P = (4\gamma\cos\theta)/D$$

Where $\gamma$ is the liquid–gas interfacial tension, $\theta$ is the contact angle on the fiber surface, and D is the effective pore diameter between adjacent fibers. According to this relation, both increasing wettability (decreasing $\theta$) and decreasing pore size (smaller D) contribute to a higher capillary pressure. Thus, a net upward driving force is established across the membrane.

At the interface between the TPU-PSA and TPU-PVP-PSA layers, the water droplet experiences two Laplace pressures, $P_1$ and $P_2$, acting in the same direction. The resulting pressure gradient $\Delta P_1$ is:

$$\Delta P_1 = (4\gamma\cos\theta_2)/D_2 - (4\gamma\cos\theta_1)/D_1$$

Where $\theta_1$ and $\theta_2$ are the contact angles of the TPU-PSA and TPU-PVP-PSA layers, respectively, and $D_1$ and $D_2$ are their corresponding pore sizes. It produces a net capillary force $CF_1$ that pulls the liquid front from the TPU–PSA layer into the TPU–PVP–PSA layer. As the liquid further approaches the TPU–PVP–PSA and HPAN interface, an additional pressure difference develops. In the schematic, $P_3$ denotes the

Laplace pressure associated with the meniscus curvature within the upper region of the TPU–PVP–PSA layer, while $P_4$ denotes the stronger capillary suction generated in the finer and more hydrophilic HPAN pore network. The second driving term is expressed as:

$$\Delta P_2 = P_4 - |P_3|$$

which contributes an additional capillary force $CF_2$ that continuously draws the liquid into the HPAN layer, enabling rapid spreading and outward transport once the hydrophilic surface is reached.

In parallel with the Laplace-pressure gradients, the stepwise increase in surface energy from the hydrophobic TPU–PSA layer to the wettability-transition TPU–PVP–PSA layer and then to the superhydrophilic HPAN layer generates a wettability-gradient driving force $F_w$. This force originates from the surface-tension imbalance along the advancing contact line and acts in the direction of decreasing contact angle, assisting the liquid front in crossing interlayer boundaries and sustaining one-way transport through the laminate.

Together, the staged capillary pressure differences $\Delta P_1$ and $\Delta P_2$, aided by the surface-energy-gradient force $F_w$, provide a passive driving mechanism that lifts fluid from the TPU–PSA side toward the HPAN side without relying on enclosed microchannels or discrete microfluidic routing. In the reverse direction, the hydrophobic TPU–PSA layer and its larger pores reduce spontaneous wetting and suppress back-penetration, helping to limit contaminant ingress from the external environment.

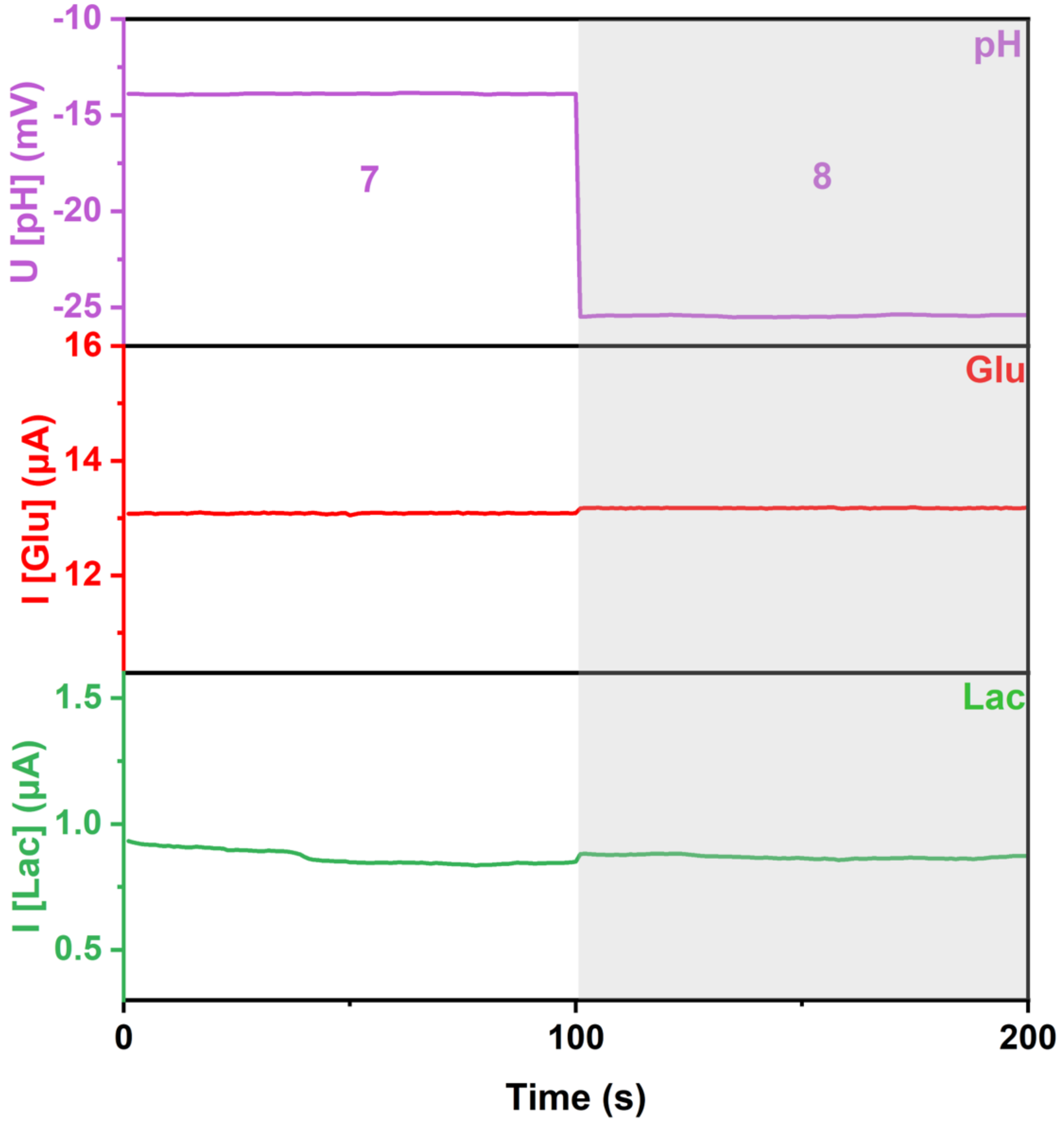


**Figure S10.** The influence of pH on the responses of sensors. Test solutions included pH 7.0 and 8.0 standard buffers with 10 mM glucose and 1 mM lactate.

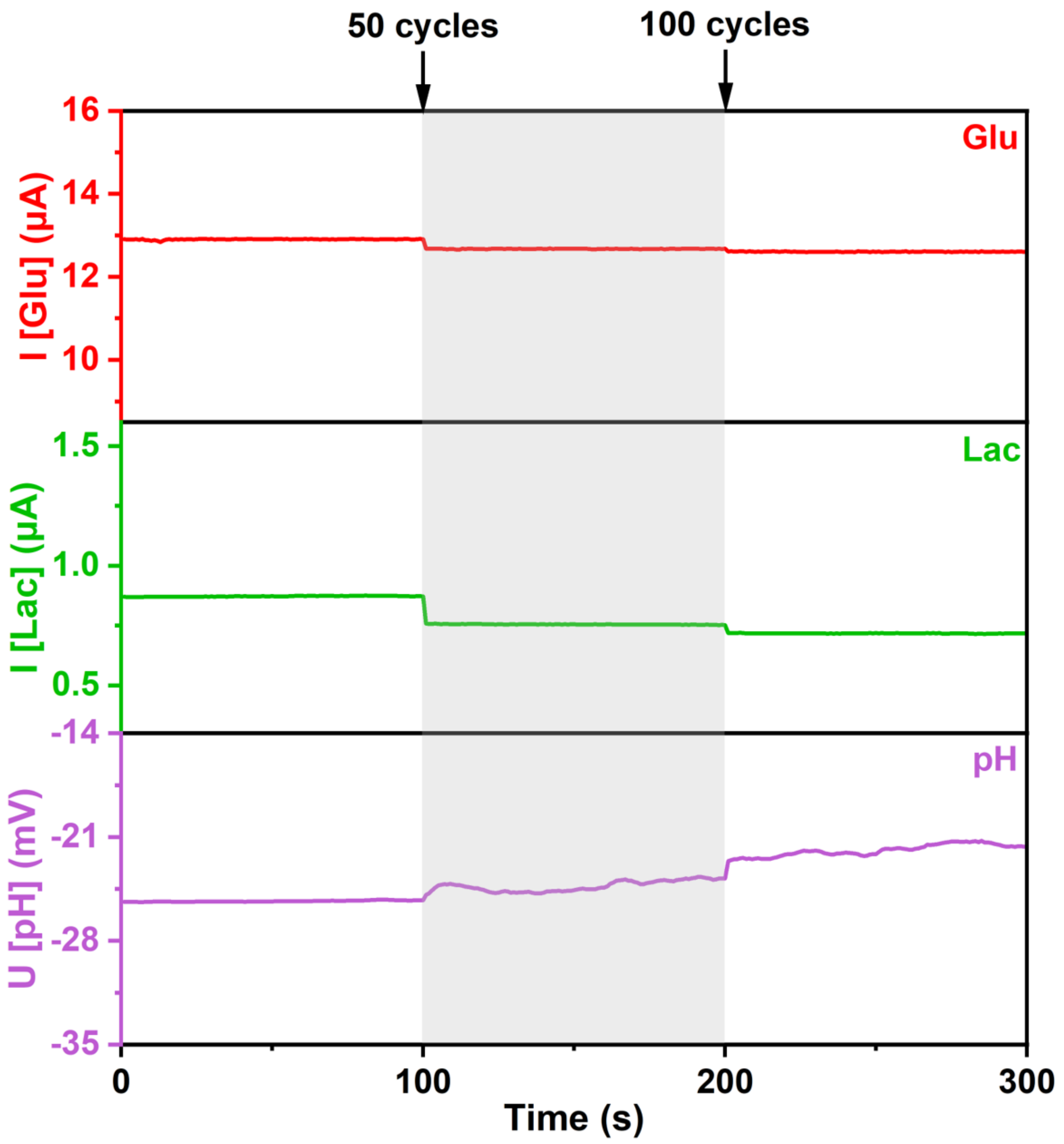


**Figure S11.** The operation stability of the sensors. Data was recorded before and after 50 and 100 cycles of repetitive operation. Test solutions included 10 mM glucose solution, 1 mM lactate solution and pH 8.0 standard buffer.

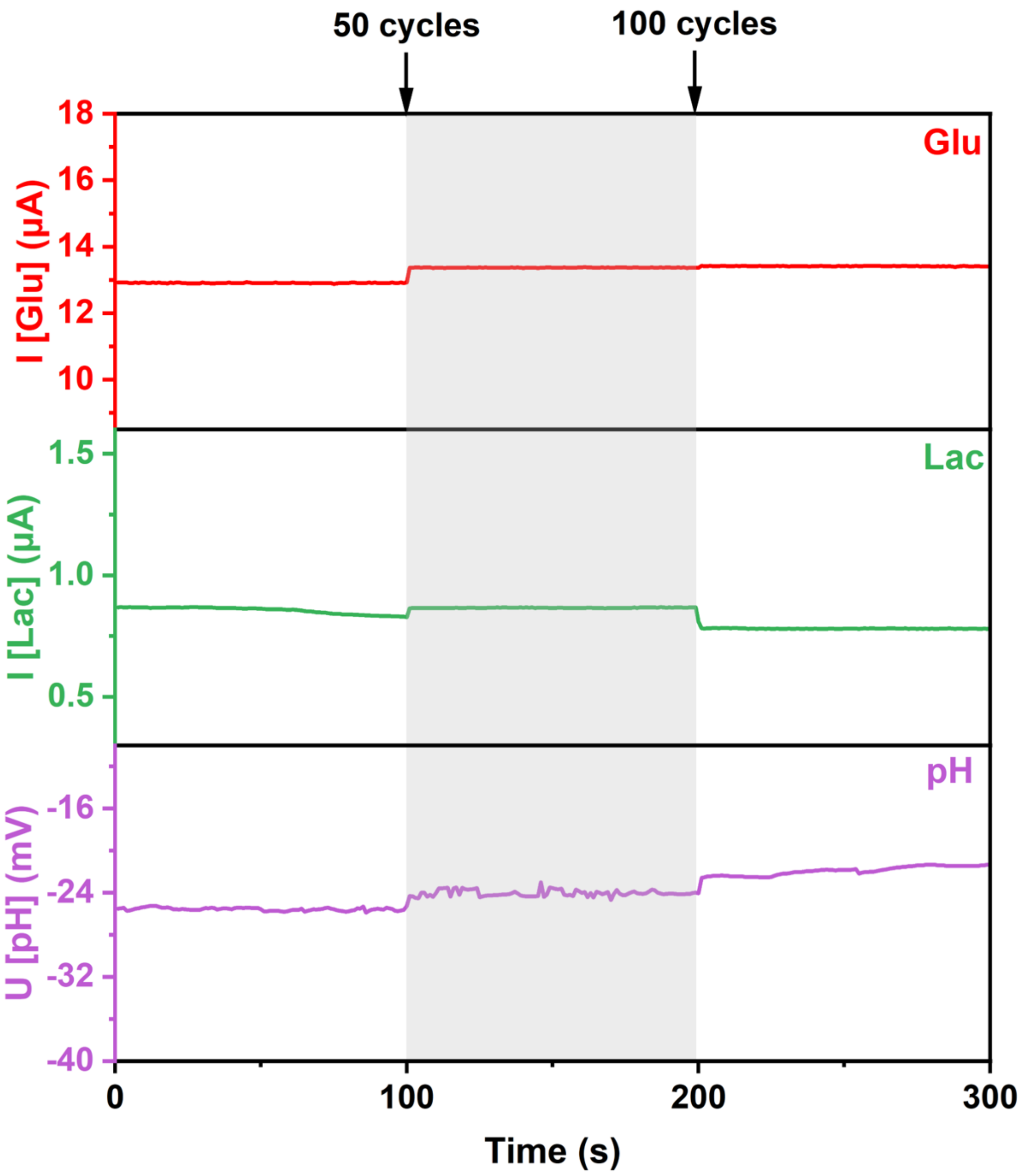


**Figure S12.** The mechanical stability of the sensors. Data was recorded before and after 50 and 100 cycles of repetitive mechanical bending (radius of curvature, 1 cm). Test solutions included 10 mM glucose solution, 1 mM lactate solution and pH 8.0 standard buffer.

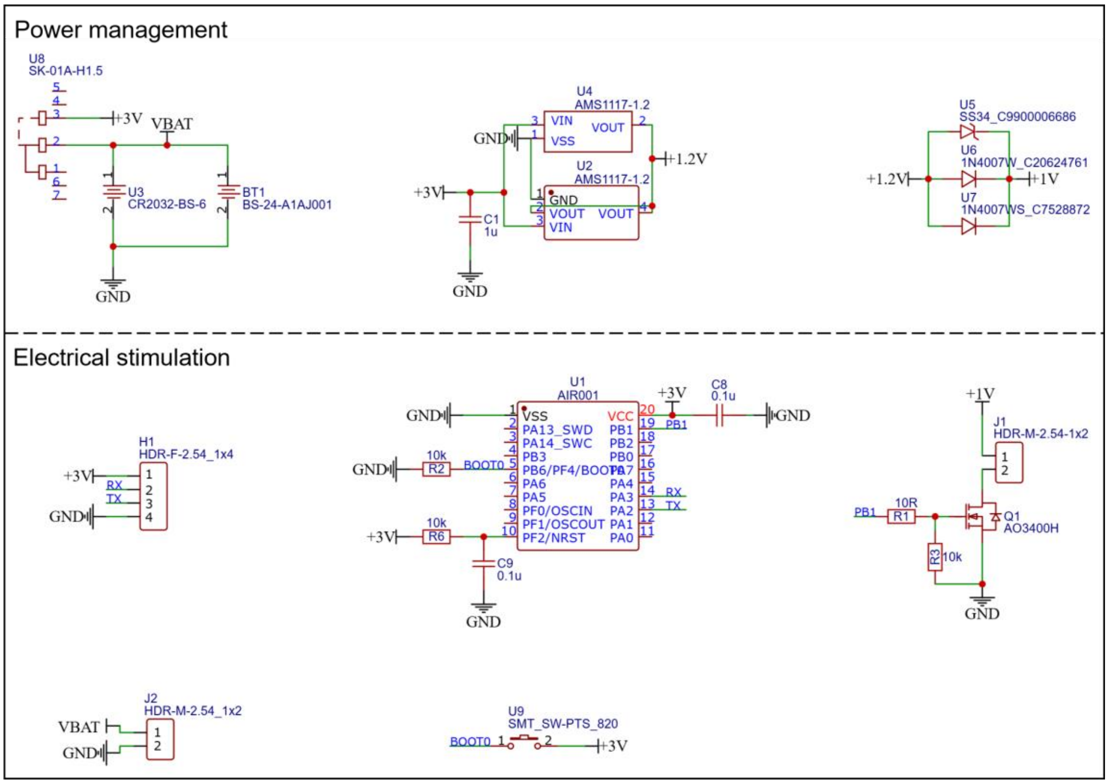


**Figure S13.** Circuit diagram of the electrical stimulation system.

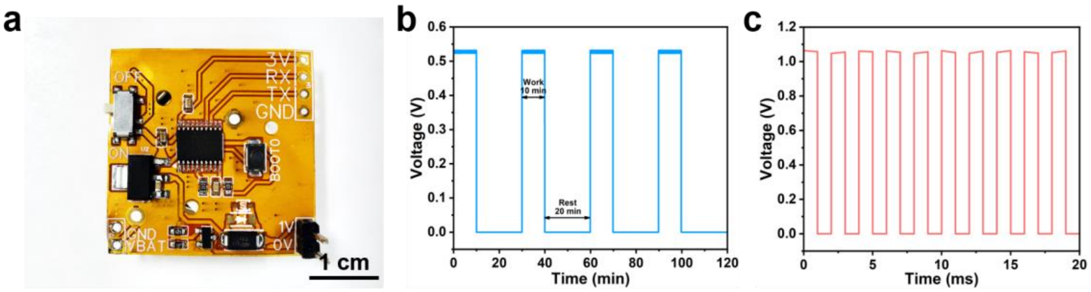


**Figure S14.** (a) Optical image of the electrical stimulation system integrated on a flexible printed circuit board (FPCB). (b) Voltage profile showing the programmed working mode of the system, consisting of a 10-minute stimulation phase followed by a 20-minute rest period. (c) Representative stimulation waveform during the working phase, delivering a 1 V, 50 Hz signal.

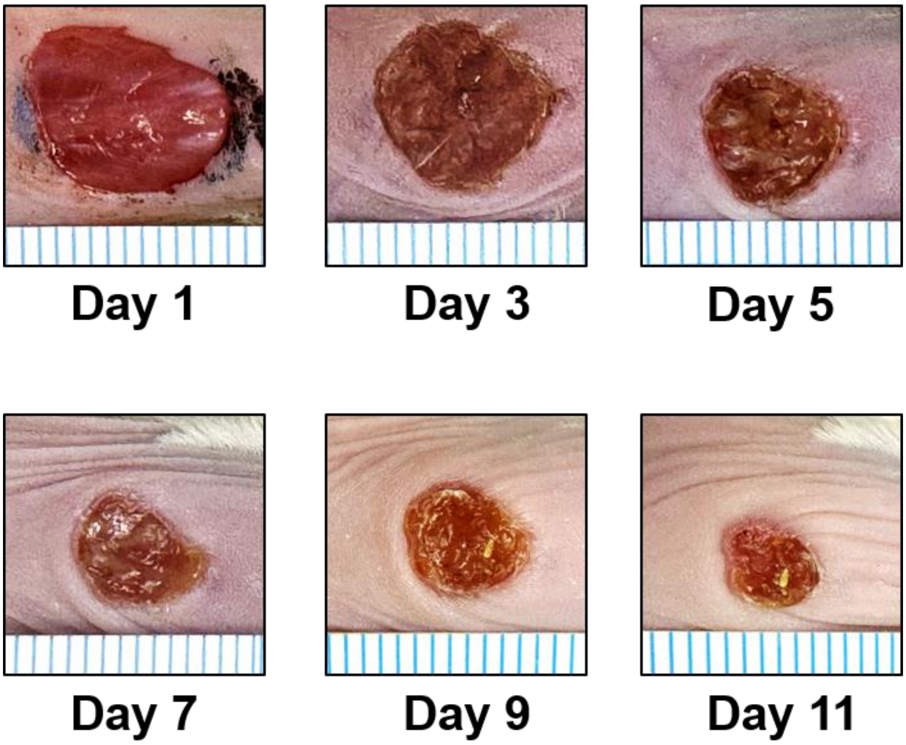


**Figure S15.** Representative photographs showing the wound healing progression over 11 days in the control group.

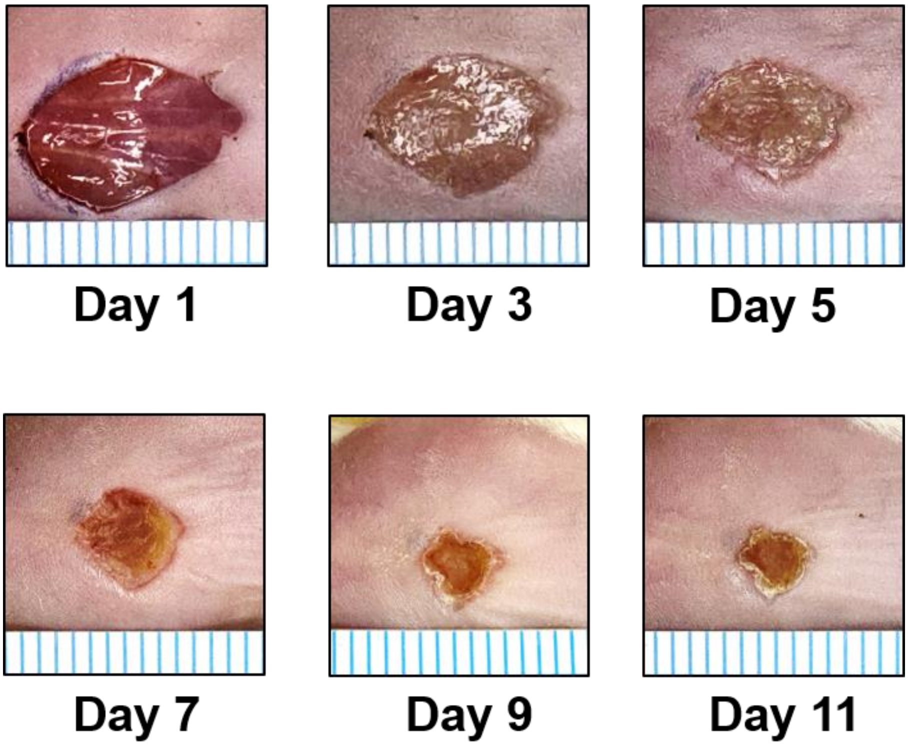


**Figure S16.** Representative photographs showing the wound healing progression over 11 days in the drug-only dressing group.

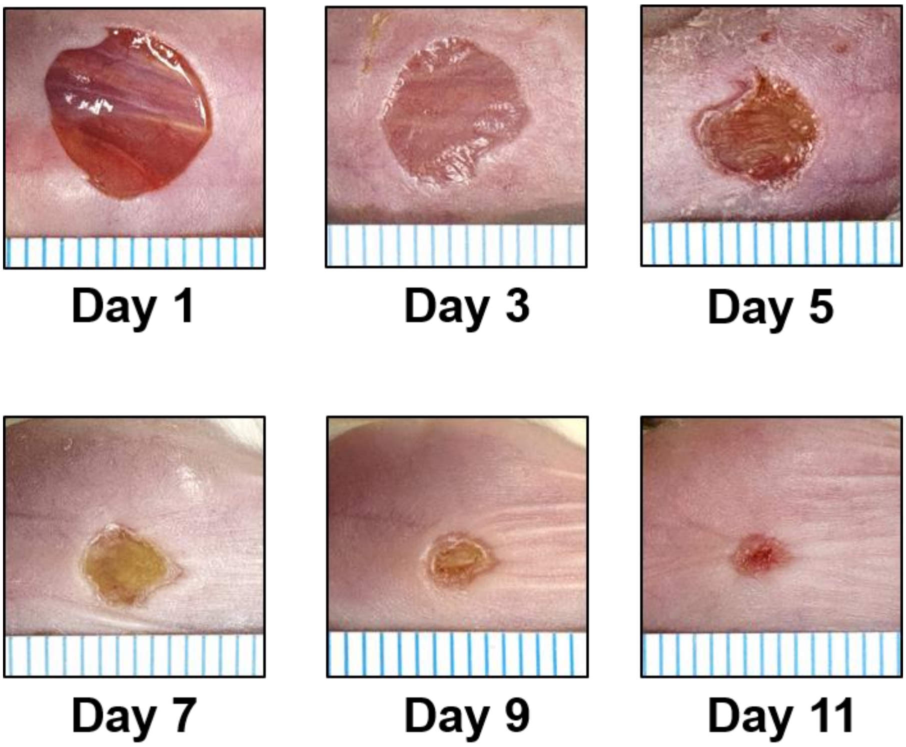


**Figure S17.** Representative photographs showing the wound healing progression over 11 days in the drug & ES dressing group.

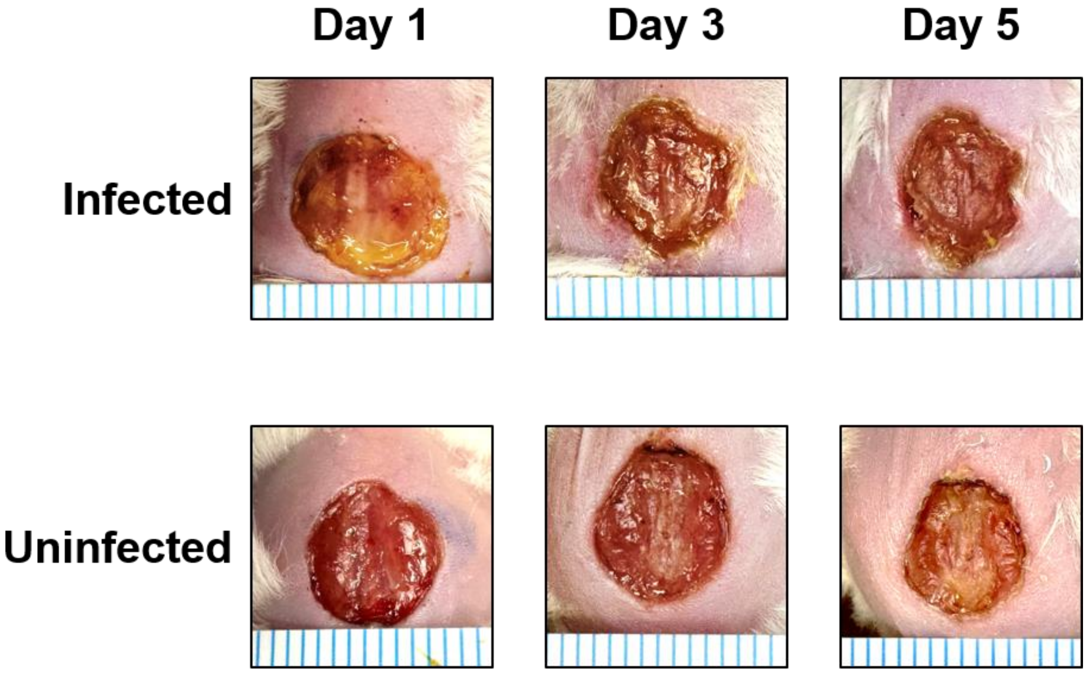


**Figure S18.** Representative photographs showing the wound healing progression at days 1, 3, and 5 in the infected and uninfected groups.

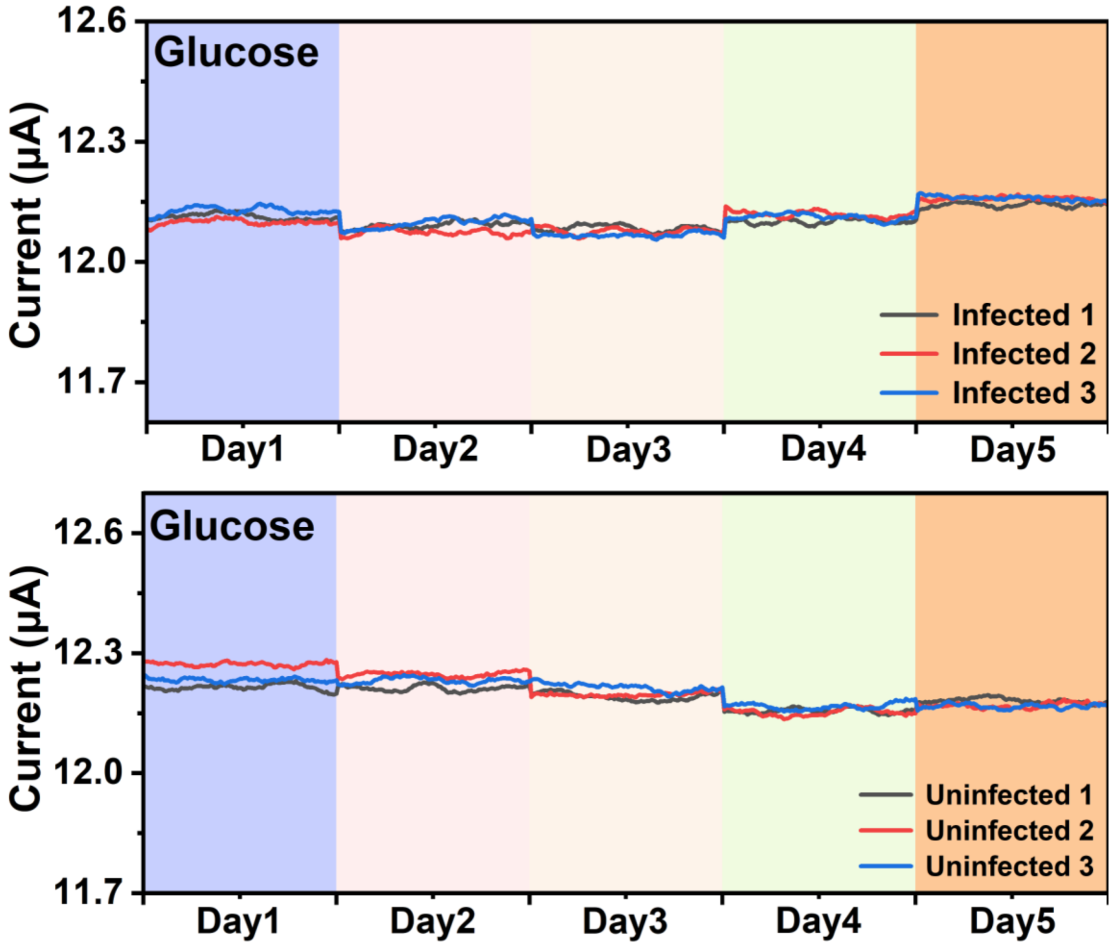


**Figure S19.** Glucose sensor signal changes over 5 consecutive days in the infected and uninfected groups under resting conditions.

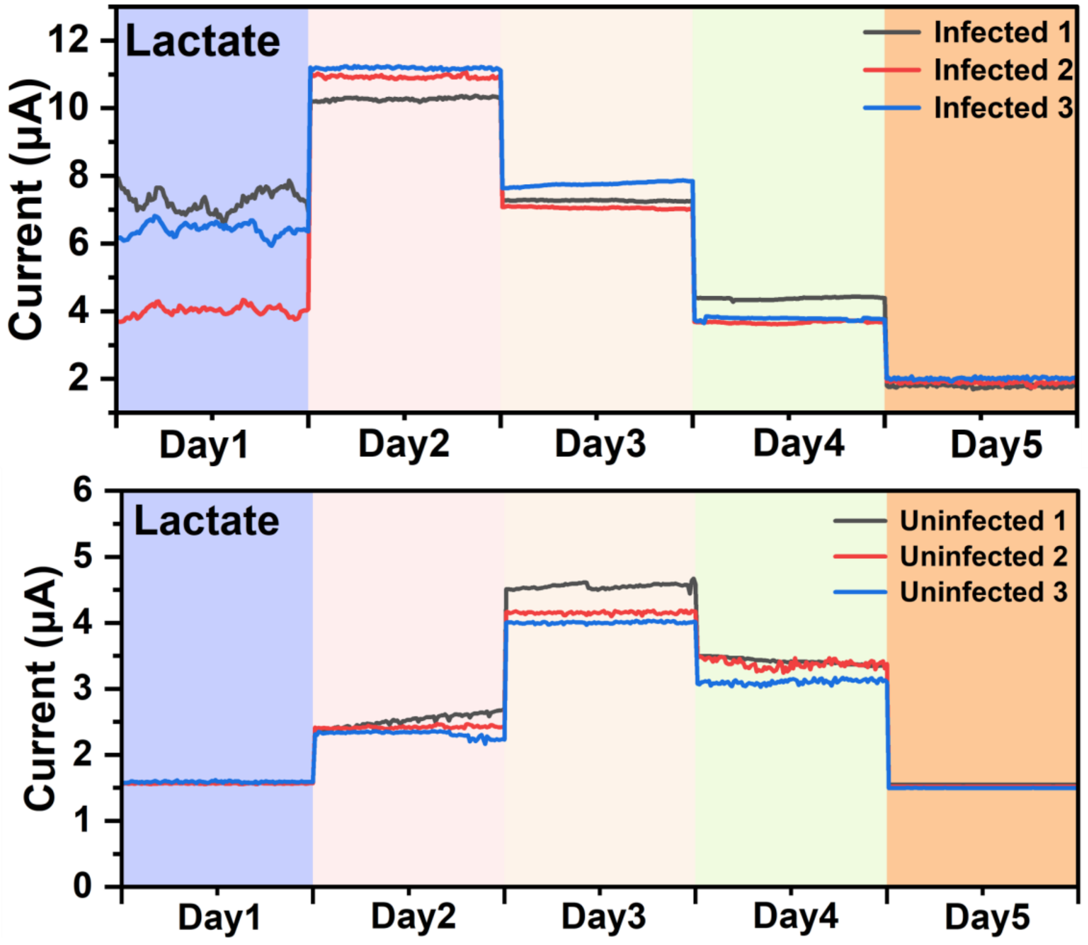


**Figure S20.** Lactate sensor signal changes over 5 consecutive days in the infected and uninfected groups under resting conditions.

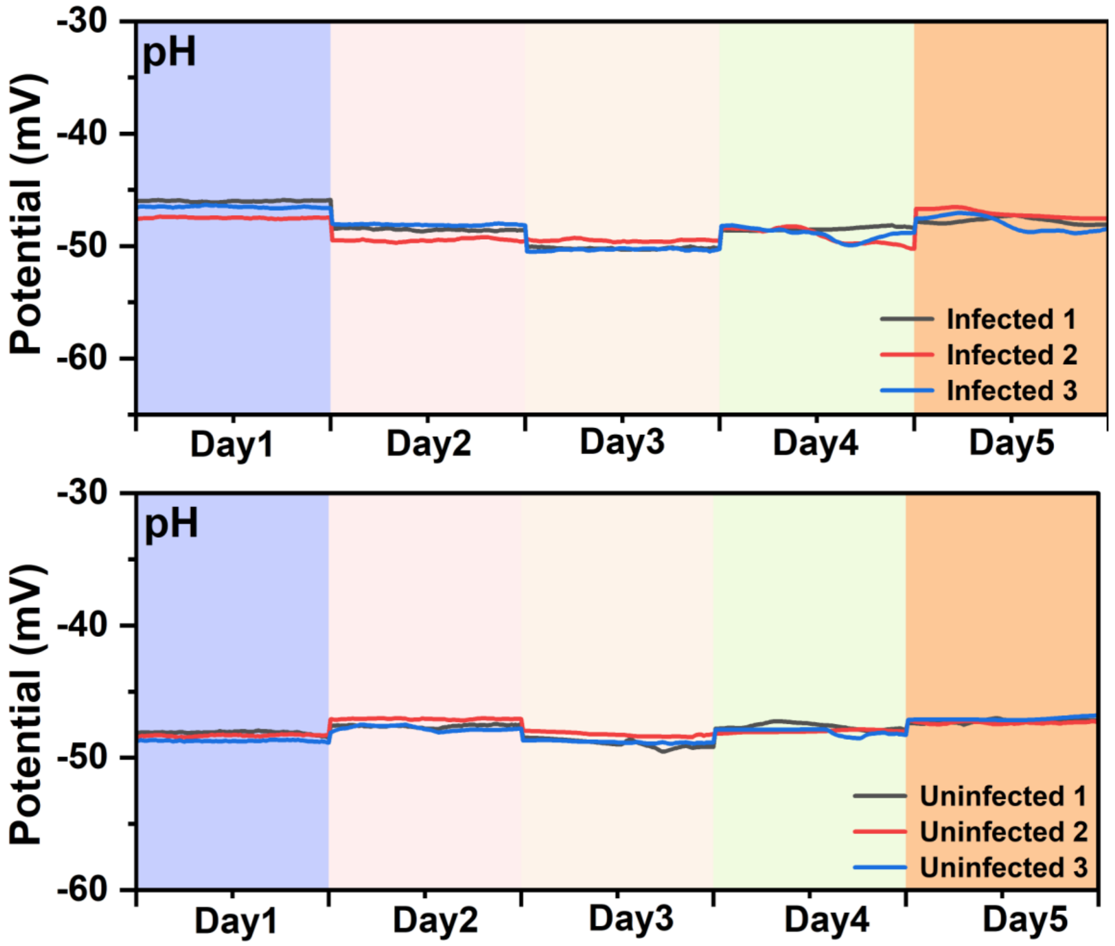


**Figure S21.** pH sensor signal changes over 5 consecutive days in the infected and uninfected groups under resting conditions.
